# Unprecedented Protein Divergence within a T3SS Family

**DOI:** 10.1101/2021.03.31.437910

**Authors:** Azzeldin Madkour, Bian Almessiry, Bertha González-Pedrajo, Brendan Kenny

## Abstract

Selection pressure drives rapid emergence of antibiotic resistance mechanisms promoting searches for therapeutic targets in bacterial processes needed for virulence, not viability, which include the Type Three Secretion System (T3SS). Distinct T3SS families evolved from the flagellar export apparatus where remaining homology hinders development of anti-T3SS specific therapies. Around 15 proteins that are highly-conserved within, but not between T3SS families yet such divergence is rarely leveraged, to promote understanding, due to unknown evolutionary histories. Here we document unprecedented divergence in two ‘LEE’ T3SS family members. Interchangeability studies uncover unusual LEE biology (eg *2-orf* genes) and illustrate each T3SS protein can tolerate dramatic change. Functional defects (12 proteins) and novel phenotypes enabled studies that reveal i) pathotype-specific protein functionality, ii) T3SS crosstalk with other processes, and iii) potential therapeutic targets. The work provides resources and testable predictions for further discoveries and will promote comparable studies between distinct T3SS families.

## Introduction

The action of antibiotics that target physiologically important bacterial processes such as flagellar movement or cell wall biosynthesis are rapidly undermined by the evolution of resistance mechanisms (Lyons and Strynadka, 2019; Roope et al., 2019). As stated by the World Health Organisation (2021) “Antibiotic resistance is one of the biggest threats to global health, food security, and development today.”. There is an emerging interest in identifying therapeutic targets against processes important for virulence, but not viability, such as Type Three Secretion Systems; T3SSs (Lyons and Strynadka, 2019; Marshall and Finlay, 2014). Transfer, via T3SSs, of ‘effector’ proteins into target cells drives physiology-altering changes underpinning many notorious diseases of man, animals, and plants (Coburn et al., 2007; Kenny and Valdivia, 2009). This ‘injectisome’ organelle evolved from the flagellar export system where significant homology hinders the development of anti-T3SS specific therapeutics (Lyons and Strynadka, 2019).

A model T3SS is provided by the Attaching and Effacing (A/E) family of enteric pathogens comprising *Citrobacter rodentium, E*.*albertii*, enteropathogenic- (EPEC) and enterohemorrhagic- (EHEC) *E*.*coli* strains (Bhatt et al., 2019; Gaytan et al., 2016; Lyons and Strynadka, 2019). This injectisome (Figure 1A) is encoded on a, horizontally-acquired, region called LEE (Locus of Enterocyte Effacement) alongside genes for transcriptional regulators, a muramidase (EtgA), protein of unknown function (Rorf1), chaperones (aid T3SS substrate stability/export process), effectors and Intimin surface protein (Bhatt et al., 2019; Gaytan et al., 2016; Lyons and Strynadka, 2019). High conservation (>80% identity) between A/E pathogen T3SS protein homologs is considered to reflect functional constraints while the more divergent nature of extracellular components (>55% identity) - translocators, effectors, Intimin - is associated with exposure to host defense mechanisms (Bhatt et al., 2019; Castillo et al., 2005; Perna et al., 1998).

**Figure 1:**
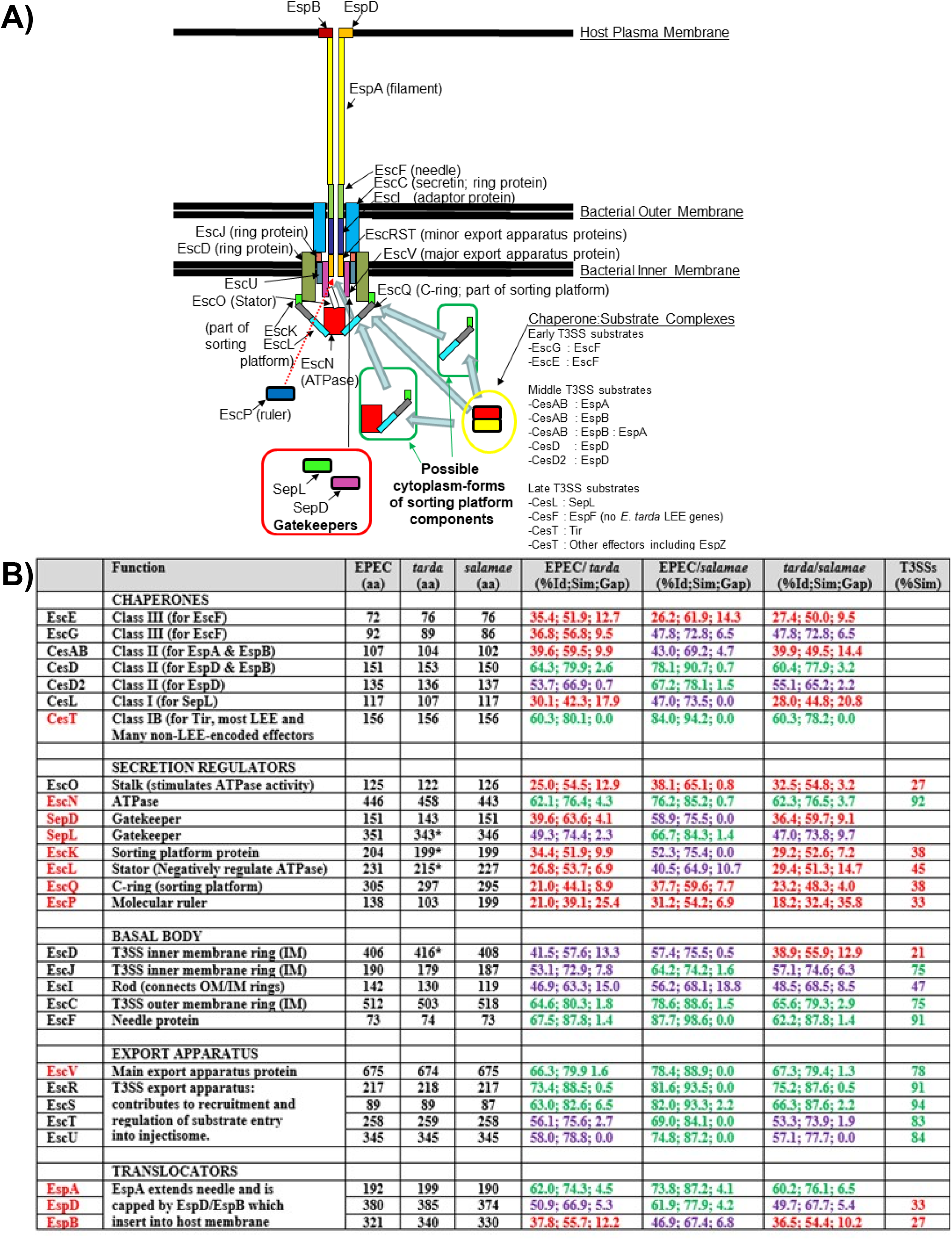
T3SS Schematic and homolog comparison data. A) Schematic of EPEC E2348/69 (EPEC) T3SS composed of a basal body: 3 ring-like structures spanning inner/outer membranes [EscD/J/C] with a cavity enclosing the export apparatus (5 proteins [EscR-V] complex providing export ‘entry’ channel) connected, via adaptor protein [EscI], to hollow needle-like structure [EscF] protruding from the cell, ii) translocon: one translocator [EspA] extends the needle and ‘capped’ by 2 pore-forming translocators [EspB/D] whose insertion into the host membrane complete the effector-delivery conduit, and iii) ATPase-gatekeeper-sorting platform complex: 8 [EscQ/K/L/N/O/P & SepD/L] proteins beneath entry channel control the order/timing of substrate export. Also indicate are cytoplasmic chaperones/substrate complexes and predicted interactions. B) lists main function of T3SS proteins alongside amino acid (aa) number as well as, in bracketa, percentage identity, similarity, gap (reflects inserts/deletions) respectively when comparing indicated EPEC, *Edwardsiella tarda* (*tarda)* or *Salmonella salamae* (*salamae)* homologs. T3SSs (%Sim) column shows reported (Diepold and Wagner, 2014) percentage similarity between homologs from multiple T3SS families. Colour coding: i) protein name in red highlights those proteins whose *tarda* homologue had no or little ability to functionally substitute its EPEC variant, ii) (%Id;Sim;Gap) column: red, purple and green lettering highlights % identity is in the very low (21%-40%), intermediate (>40%-60%) or high (>61-74%) category, respectively.

Most knowledge on T3SS protein functionality relates to studies on prototypic systems from 5 families where homologs share as little as 21% similarity (Diepold and Wagner, 2014). Such divergence argues for selection pressure driving changes beneficial to the host bacteria’s lifecycle leading to a pathotype-specific T3SS. While such divergence provides opportunities for interrogating T3SS protein biology and developing anti-T3SS (broad- or pathotype-specific) therapeutics, it is rarely leveraged due to unknown evolutionary histories (Klein et al., 2017).

Here we describe how our interest in an apparently degenerate LEE region led to the uncovering of two, overlooked, examples of unprecedented divergence within an evolutionary closely-related T3SS family. We reveal this region is not degenerate but has unusual biology while studies with the homologs i) undermine the idea of high LEE T3SS protein conservation reflecting functional constraints, ii) illustrate many, if not all the T3SS proteins possess pathotype-specific functionality, iii) reveal T3SS crosstalk with other processes, iv) provide a strategy for identifying T3SS- and/or pathotype-specific therapeutic targets, v) forward testable predictions and vi) describe phenotypes and tools to enable future discoveries.

## RESULTS

### Unprecedented protein divergence within the LEE T3SS family

The report of an invasive (non-A/E) fish pathogen, *Edwardsiella tarda* FPC503 (*tarda*), carrying a LEE region lacking all but one A/E pathogen-conserved effector (Nakamura et al., 2013) indicated repurposing for *tarda*-specific functionality. However, the apparent absence/truncation of many genes (∼32%) prompted data re-interrogation revealed sequences for all but 8 (*cesF, rorf1, grlA, grlR, map, espF*-*H*; see Figure 1A); latter dispensable for A/E pathogenesis and T3SS functionality (Deng et al., 2004; Ruano-Gallego et al., 2015). Moreover, the analysis revealed a second effector (EspZ) and a 2-*orf eae* (encodes Intimin) gene while 4 genes were not truncated, but disrupted by sequencing-verified: 14bp insertion (*escD*), stop codon (*escK*) and frameshift (*sepL, escL*) mutation. However, 2-*orf* genes are expressible by, for example, ribosome-mediated mechanisms (Weiss, 1991). Unexpectedly, comparing the deduced protein sequences to their EPEC (E2348/69) counterpart revealed undocumented high divergence between LEE homologs: 21%-73.4% identity (Figure 1B).

### A majority of *tarda* LEE T3SS proteins functionally replace their EPEC counterparts

Such unprecedented divergence prompted assessment of whether the *tarda* genes could functionally substitute their EPEC counterpart. Thus, *tarda* genes were PCR-cloned and introduced into the appropriate EPEC T3SS gene-deficient strain to assess T3SS functionality using two well-established assays (Kenny et al., 1997a; Kenny et al., 1997b). Strain genotype was routinely supported by PCR analysis (not shown).

These studies revealed 16 (of 28) genes functionally substituted, as exemplified by *cesAB* that encodes a chaperone 39.6% identical to EPEC CesAB (Figure 2A). EPEC Δ*cesAB* secretes only one of three translocator proteins (Figure 2B) while Tir, the first and most abundantly-delivered effector (Mills et al., 2008), is not transferred into mammalian (HeLa) cells (Figure 2C). These defects were rescued by reintroducing EPEC or *tarda cesAB* (Figure 2B and 2C). Thus, despite high divergence (Figure 2A) *tarda* CesAB functionally replaces EPEC CesA/B.

**Figure 2:**
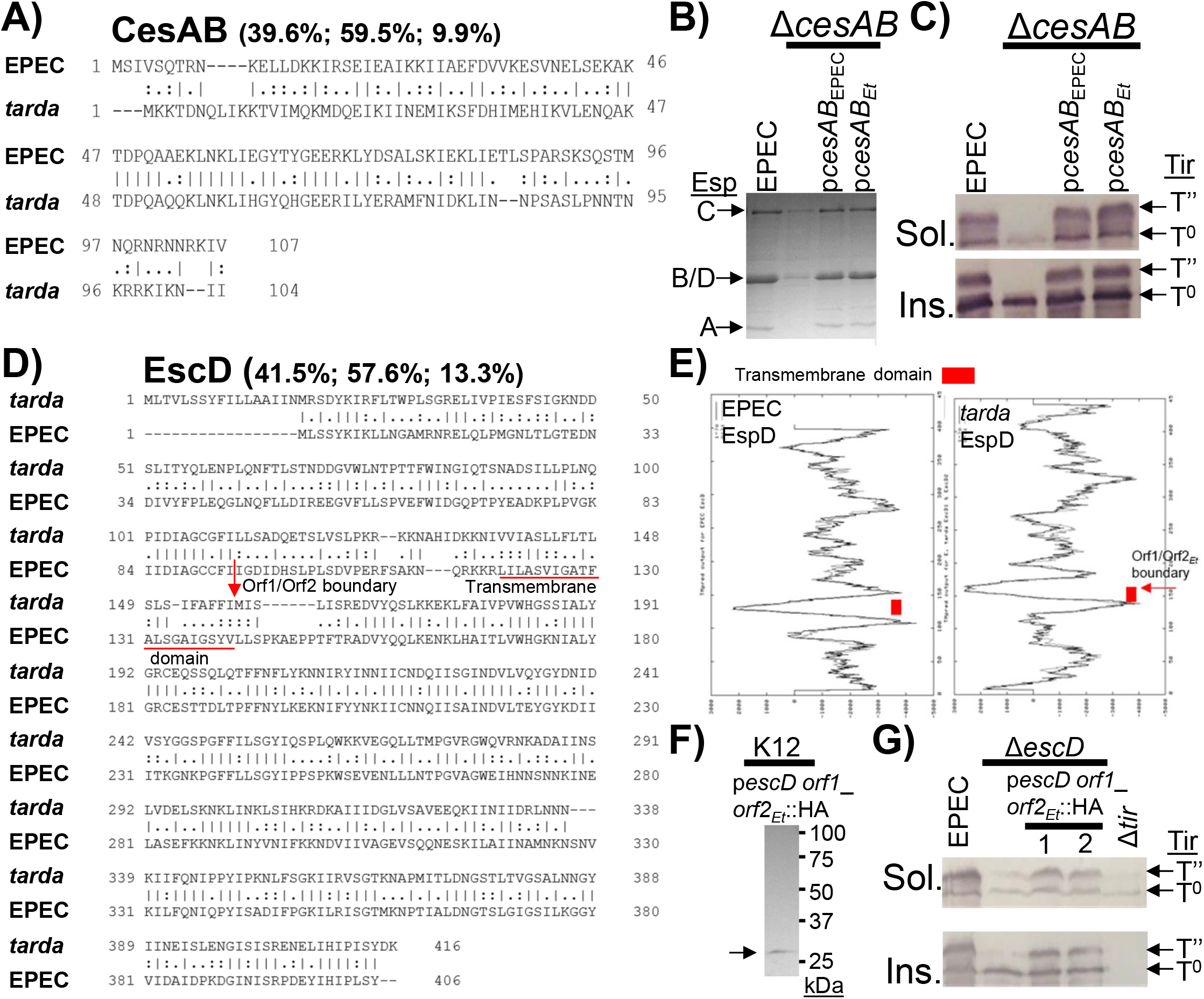
Examples of functionally substituting *E. tarda* genes/proteins. Sequence alignment of EPEC and *E. tarda* (*tarda)* A) CesAB (chaperone) and D) EscD (basal body) homologs. Indicated, within brackets, are % identity/similarity/gap scores, respectively with residues either identical (vertical line), similar (double dot), different (single dot) and absent (dashed line) residues. Location of predicted *tarda* EscD Orf1/Orf2 boundary and transmembrane segment are shown (D, E). Indicated strains were used to infect tissue culture media (B), HeLa cells (C, G) and lysogenic broth (F) before isolating secreted proteins (B), Triton X-100 soluble (Sol.; contains host cytoplasm and membrane proteins plus T3SS-delivered effectors) and insoluble (Ins.; contains host nucleus and cytoskeleton plus adherent bacteria) fractions (C, G) and total bacterial extracts (F). Samples were resolved (SDS-PA gels) and proteins visualised by Coomassie blue staining (B) or Western blot analysis; latter detecting Tir effector (C, G) or HA-tagged proteins (F). Arrows indicate position of secreted translocator (EspA, EspB, EspD) and autotransporter (EspC) proteins (B), unmodified (T^0^) and host kinase-phosphorylated (T’’) Tir forms (C, G) and HA-tagged protein (F). Tir-Intimin interaction is evidenced by Tir T’’ in the Insol. fraction (Kenny et al., 1997b; Kenny and Finlay, 1997). Strains were non-pathogenic (K12) *E*.*coli*, EPEC, Δ*cesAB* (lacks chaperone) and Δ*escD* (lacks inner membrane ring protein) with or without indicated plasmids (*Et* denotes from *E. tarda*).

Notably one substituting gene was interrupted (14bp insertion) separating the encoded N-terminal (carries sole predicted sec pathway export signal and membrane-spanning region) and C-terminal (periplasmic) domains (Figures 2D, 2E and S1_i). It was predicted that *tarda* EscD would, as per EPEC EscD, produce a single full length polypeptide. However, adding an epitope tag (human influenza hemagglutinin; HA) at the Orf2 C-terminus revealed expression of a single Orf2-sized (∼28kDa), not Orf1 (∼14kDa) or full length (∼42kDa), fusion protein. Disrupting *orf1* (not shown) compromised the export process unlike HA-tagging Orf2 (Figure 2G) supporting independent Orf1/Orf2 expression. This work illustrates that *tarda* LEE *2-orf* genes can produce functional proteins but perhaps not via predict ribosome-mediated mechanisms.

The 16 ‘functionally-substituting’ members are 9 (of 10) basal body-export apparatus components, 6 (of 7) chaperones and 1 (of 8) ATPase-gatekeeper-sorting platform complex proteins (Figure 1B, 2 and S1_i-xv). Functional interchangeability was not correlated with sequence divergence as members were distributed in all designated homology (% identity) category: very low (21-40%; EscO [21%] <CesL <EscE <EscG <CesAB [39.6%]), intermediate (>40-60%; EscD <EscI <EscJ <CesD2 <EscT <EscU) and highest (>60-74%; EscS <CesD <EscC <EscF <EscR). Thus, many LEE T3SS proteins can accommodate substantial change without obvious impact on T3SS functionality.

### Complementation defects correlate with protein divergence

Of the remaining 12 proteins, half were clearly expressed as restored some T3SS activity; latter exemplified by another *2-orf* gene (Figure 3A) which encodes a SepL homolog (49.3% identity; Figure 3B). As reported (Deng et al., 2005), EPEC Δ*sepL* secretes, not translocator but effector proteins, most prominently Tir and NleA (Figure 3C). The mutant also fails to deliver Tir into HeLa cells with all defects rescued re-introducing EPEC *sepL* (Figure 3C and 3D). By contrast introducing *tarda sepL* only rescued one defect, partially, as enabled some Tir delivery (Figure 3C and 3D). Epitope-tagging Orf2 supported production of a full-length SepL protein (Figure 3E). The 6 ‘partial functionality’ members (Figure S2_i-v) were a chaperone (CesT), 2 secretion regulators (SepL, EscP) and all translocators (EspA, EspB, EspD) distributed among the homology categories (Figure 1B): very low (EscP <EspB), intermediate (SepL <EspD <CesT) and highest (EspA).

**Figure 3:**
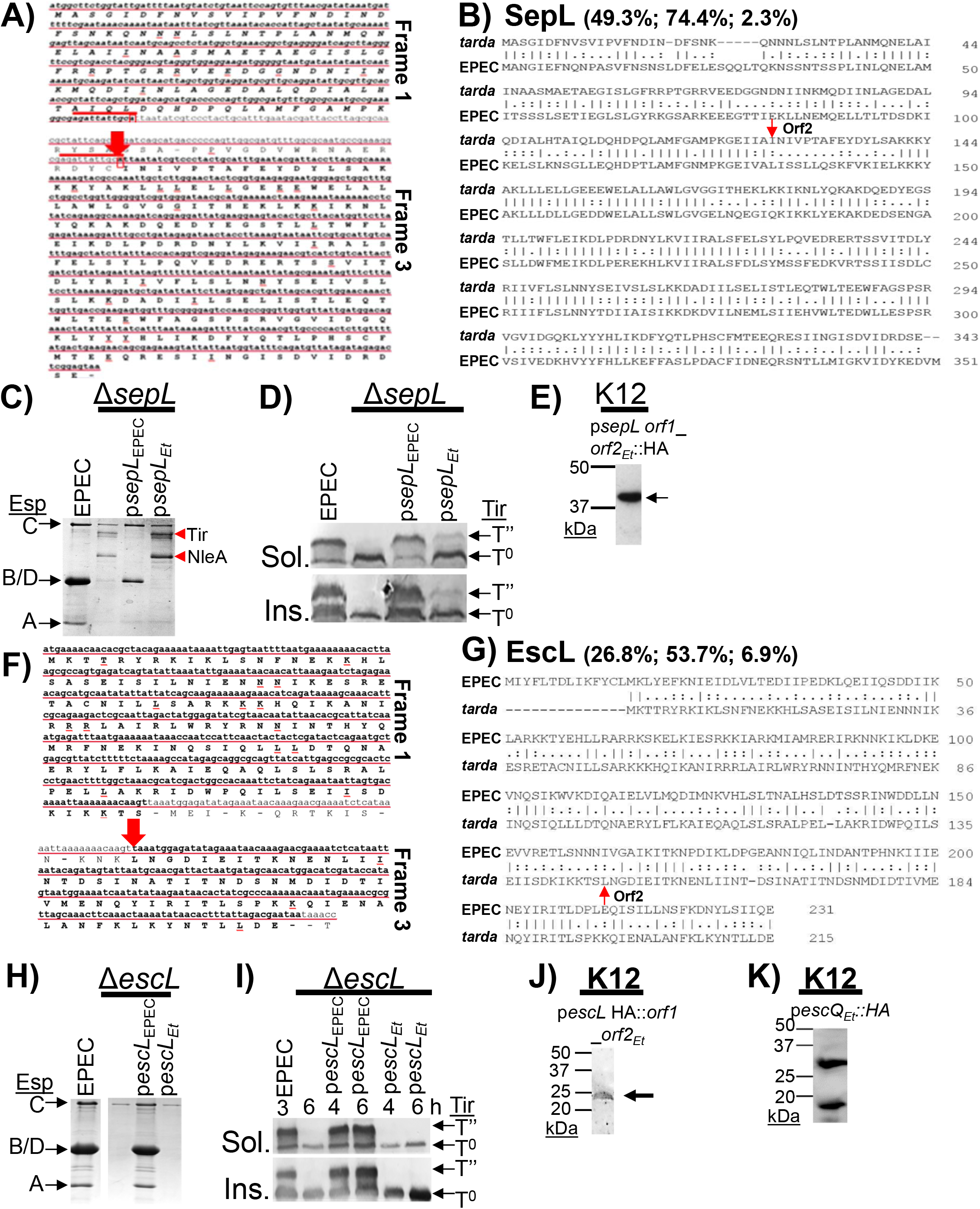
Examples of *E*.*tarda* variants displaying complementation defects. Gene sequences of 2-*orf E. tarda* (*tarda)* A) *sepL* and F) *escL* genes; position of frameshift mutation indicated (large red arrow). Sequence alignment of EPEC and *tarda* B) SepL and G) EscL homologs. Indicated strains were used to infect tissue culture media (C, H), HeLa cells (D, I) and lysogenic broth (E, J, K) prior to isolating secreted proteins (C, H), Triton X-100 soluble (Sol.) and insoluble (Ins.) fractions (D, I), or total bacterial extracts (E, J, K). Samples were resolved (SDS-PA gels) and proteins visualised by Coomassie blue staining (C, H) or Western blot; latter detecting Tir effector (D, I) or HA-tagged fusion proteins (E, J, K). Strains were non-pathogenic (K12) *E*.*coli*, EPEC, Δ*sepL* (lacks gatekeeper), Δ*escL* (lacks ATPase activity regulator) with or without indicated plasmids. Information on sequences, fraction composition, abbreviations and arrows as per Figure 2.

The final 6 *tarda* genes failed to replace their EPEC variant as exemplified by another 2-*orf* gene (Figure 3F) encoding an EscL homolog (26.8% identity; Figure 2G). Thus, the Δ*escL* mutant defect was rescued by plasmid re-introducing EPEC, not *tarda escL* (Figure 3H and 3I). HA-tagging supported expression of a full-length EspL protein (Figure 3J). Similar findings were obtained for other examined members: SepD, EscQ, EscN and, final 2-*orf* encoded protein, EscK (Figure S3_i-iv). Notably *tarda* EscQ produced full length and C-terminal polypeptides (Figure 2K) as predicted for its EPEC variant (Gaytan et al., 2016). The 6 ‘complementation-null’ members were 5 (of 8) ATPase-gatekeeper-sorting platform complex proteins and 1 (of 5) export apparatus proteins in the very low (EscQ <EscL <EscK <SepD) or highest (EscN <EscV) homology categories (Figure 1B).

The demonstration of 27 LEE *tarda* (includes four 2-*orf*) genes producing full-length and/or functional proteins suggests the complementation defects are not due to non-expression but reduced expression or, more likely, sequence divergence. To explore this hypothesis studies focused on the final ‘complementation-null’ group member, EscV as in highest homology category (66.3% identity; Figure 1B and S3_v). The EscV C-terminal domain provides the T3SS export gate and SepL docking sites (Gaytan et al., 2018; Portaliou et al., 2017). Divergence within multiple docking sites (Figure 4A) prompted exchanging the entire C-terminal region for that from EPEC generating a chimera that rescued all export defects (Figure 4B and 4C) clearly linking the defects to divergence.

**Figure 4:**
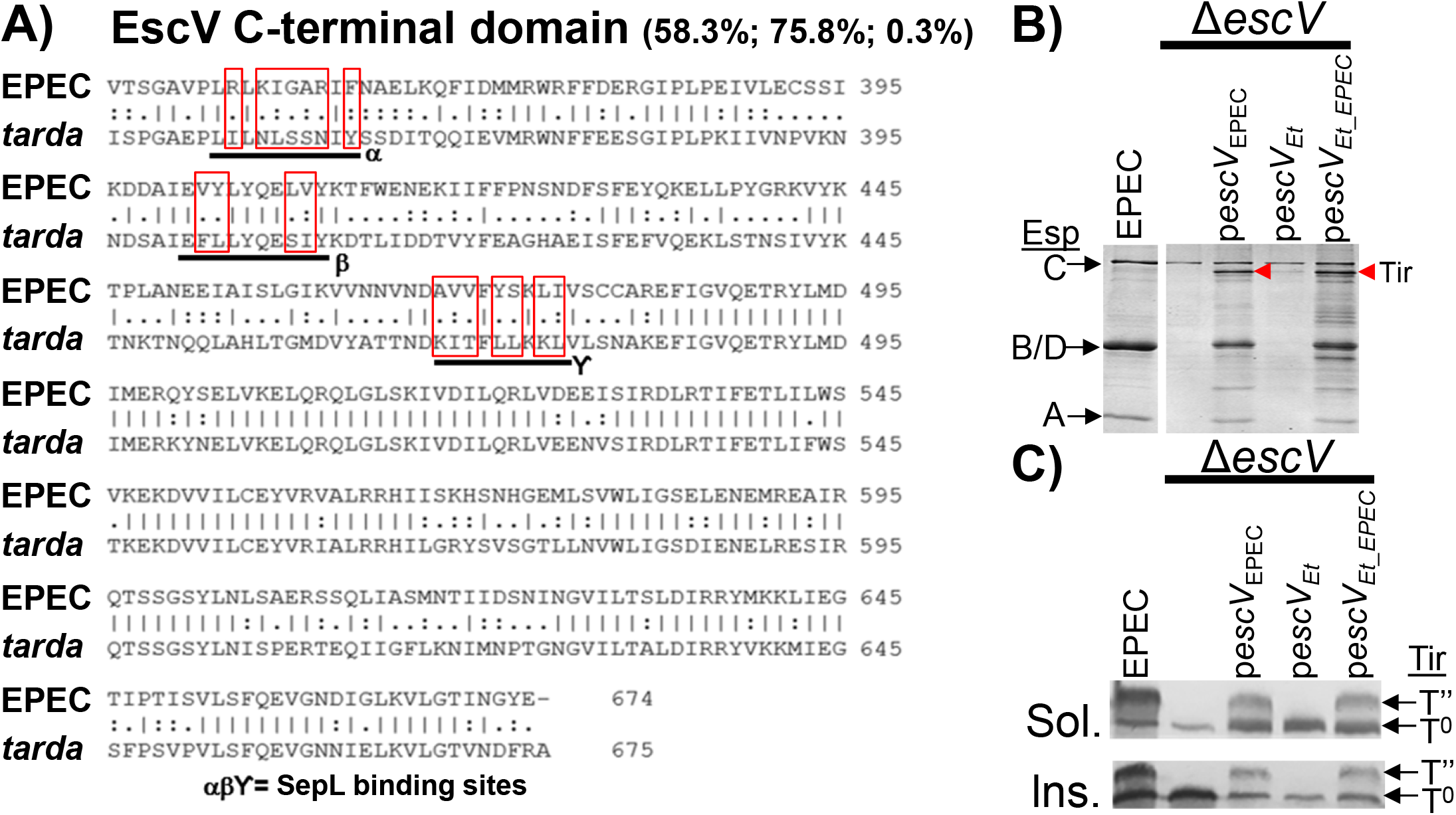
Complementation defects-protein divergence linkage. Alignment of EscV C-terminal domain from EPEC and *E. tarda* (*tarda)* with red rectangle highlighting divergence in SepL (gatekeeper) binding sites (Portaliou et al., 2017). Indicated strains were used to infect tissue culture media (B) and HeLa cells (C) before isolating secreted proteins (B) and Triton X-100 soluble (Sol.) and insoluble (Ins.) fractions (C). Samples were resolved (SDS-PA gels) and protein visualised by Coomassie blue staining (B) or Western blot to detect the Tir effector (C). Strains used were EPEC, Δ*escV* (lack EscV export protein) with or without indicated plasmids; p*escV*_*Et*_EPEC_ encodes a chimera protein; *E. tarda* 1-334; EPEC 345-674. Information on sequences, fraction composition, abbreviations and arrows as per Figure 2

### Anti-T3SS therapeutic targets

Next we examined if complementation defects could be leveraged to define protein features or residues important for T3SS functionality to provide potential therapeutic targets. This work focused on the partially-complementing translocator proteins (weakly restore Tir delivery; Figure 5A) as their extracellular location offer advantages for therapeutic development (Lyons and Strynadka, 2019). Notably, Tir delivered by EPEC expressing the most-divergent *tarda* translocator, EspB (37.8% identity; Figure S2_i) stably interacted with Intimin unlike Tir delivered by the *tarda* EspA- or EspD-expressing strains (Figure 5A). Rescuing these defects by co-expressing the 3 *tarda* translocators (Figure 5B and S2_i/ii) is suggestive of co-divergence i.e. selection-driven change/s that negatively impact on T3SS functionality ‘compensated’ by change/s in partner protein/s.

**Figure 5:**
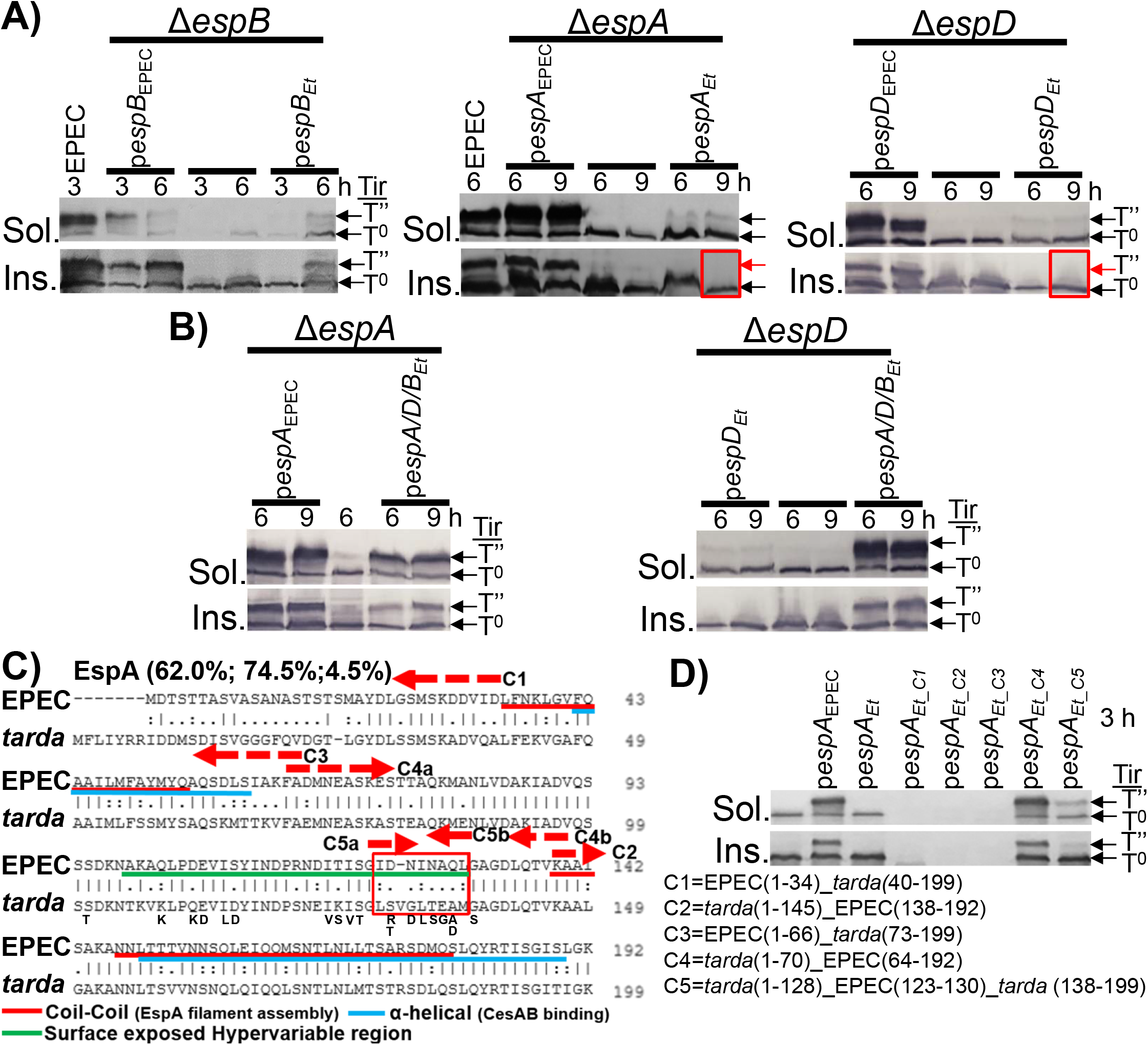
Interchangeability and domain-swap experiments reveal co-divergence and changes underpinning complementation defects. Indicated strains were used to infected HeLa cells (3h or specified times) before isolating Triton X-100 soluble (Sol.) and insoluble (Ins.) fractions for Western blot analysis probing for the Tir effector (A, B, D). Sequence alignment of EPEC and *E. tarda* (*tarda)* EspA homologs showing key features and information on generated chimera (C, D). Red rectangle highlights region where divergence is linked to complementation defect with natural variation (found in other A/E pathogen homologs) in this region indicated below the *tarda* sequence. Strains used were EPEC and strains lacking EspB (Δ*espB)*, EspA (Δ*espA*) or EspD (Δ*espD*) translocator proteins with or without indicated plasmids. Information on sequences, fraction composition, abbreviations and arrows as per Figure 2

To define complementation-interfering changes in *tarda* EspA (60% identity; Figure 5C) four chimera were initially generated (Figure 5C). Three chimera (C1-C3) reduced the levels or apparent molecular mass of examined effectors - Tir (Figure 5D) and EspF/EspB (Figure S2_ii) - suggesting EspA may have a cryptic role in regulating, directly/indirectly, T3SS substrate expression or stability levels. The fourth chimera (C4) rescued the defects (Figure 5D) implicated divergence in the final 54 residues prompting a focus on the most divergent area: residues 128-137 (0% identity; Figure 5C). Swapping the 10 *tarda* residues for EPECs 9 generated a chimera (C5) restored significant levels of Tir delivery and Tir-Intimin interaction (Figure 5C) reporting a key role for this with other, undefined, C-terminal features.

Similar studies with *tarda* EspD (50.9% identity; Figure 6A) initially involved 2 chimera (C1, C2; Figure 6A) implicating divergence in the final 49 residues (Figure 6B); latter provides a predicted secretion signal (Deng et al., 2015) and coil-coiled feature (CCF) involved in oligomerisation and membrane-insertion events (Chatterjee et al., 2015; Daniell et al., 2001). Replacing the final *tarda* 8 residues for EPEC’s increased Tir delivery levels (Figure 6B) supporting it’s predicted ‘secretion signal’ role and linking the majority of the complementation defect to CCF region divergence. Disrupting the CCF, by substituting 3 residues (A_340_, A_347_, Q_354_) to arginine, led to phenotypes (Daniell et al., 2001) equivalent to those of the *tarda* EspD variant. Absence of phenotypes following substitution of A_340_ or A_340_A_347_ residues (Daniell et al., 2001) suggests Q_354_ plays the key role. Notably, these three residues are conserved in all 50 examined A/E pathogen EspD variants, including an unusually divergent one (EspD_multi_). By contrast the corresponding *tarda* EspD residues were A, G and L i.e. two non-conservative changes (Figure 6C). Surprisingly, the EspD_multi_ and triple-substituted variants had predicted C-terminal CCFs (Figure 6D) and, whilst absent from *tarda* EspD, could readily be generated by changing two residues to their EPEC counterpart (Figure 6E). These findings argue for Q_354_, not the CCF *per se*, being important for stable, EspD/EspA-dependent, Tir/Intimin interaction. Attempts to substitute Q_354_ to its *tarda* counterpart, L, in untagged and N-terminal epitope-tagged EspD variants were unsuccessful (changed compromised protein expression or stability; not shown) suggesting this residues has other important roles in EPEC EspD biology.

**Figure 6:**
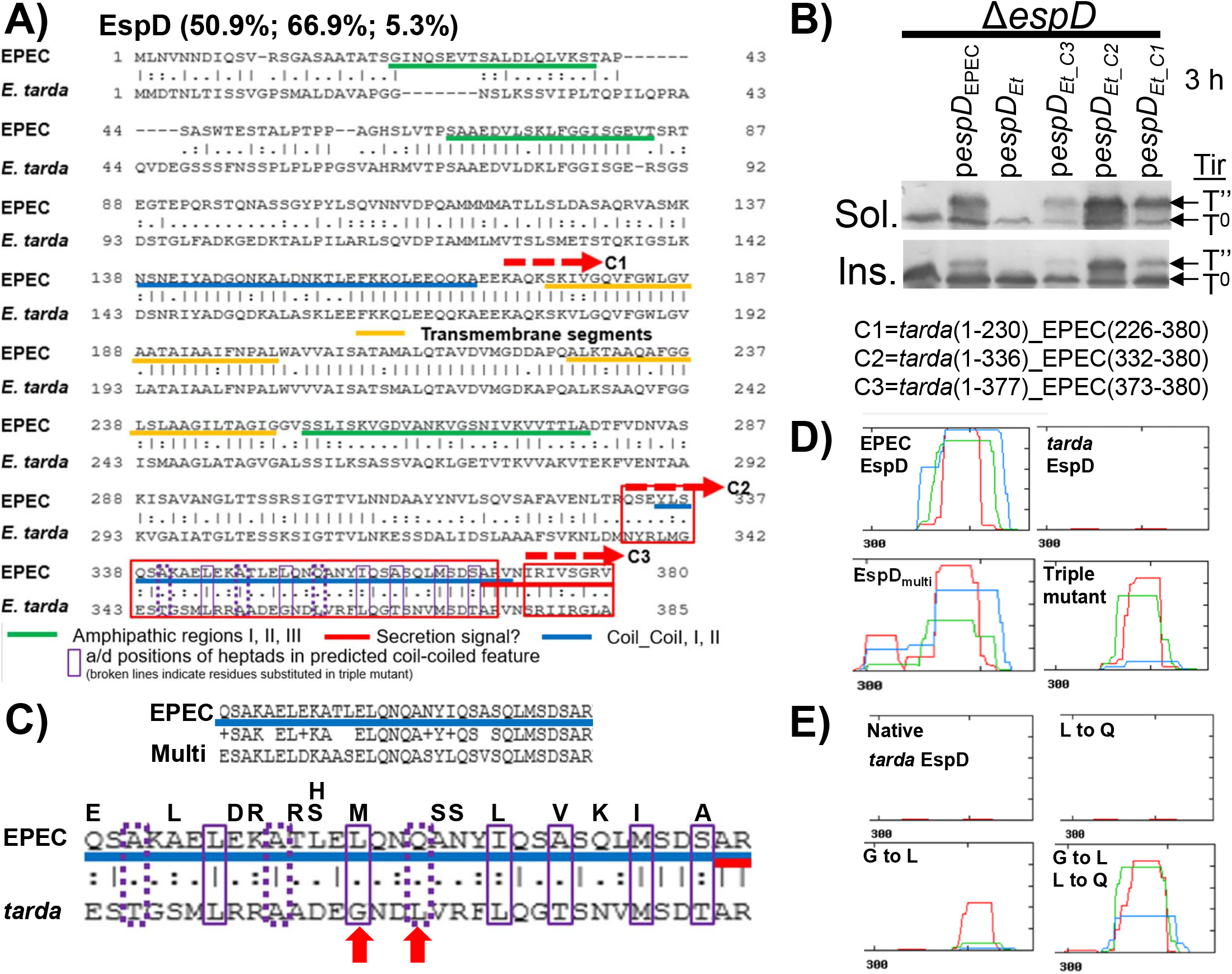
Domain swapping reveals EspD features for Tir delivery and stable Tir-Intimin interaction. A) Sequence alignment of EPEC and *E. tarda* (*tarda)* EspD homologs showing key features and information on generated chimera (A, B). Red rectangle highlights region where divergence is linked to complementation defects (A). CCF region comparisons between EPEC/*tarda* and EPEC/Multi (A/E pathogen variant with many substitutions) sequences with natural variation (found in other A/E pathogen homologs) indicated above the EPEC sequence (C). Red arrows highlight two non-conservative differences (C). CCF prediction output for indicated EspD variants (D, E). HeLa cells were infected with the Δ*espD* mutant carrying, or not, the indicated plasmid before isolating Triton X-100 soluble (Sol.) and insoluble (Ins.) fractions for Western blot analysis to detect the Tir effector (B). Information on sequences, fraction composition, abbreviations and arrows as per Figure 2

Collectively the work illustrates that i) domain-swapping experiments can identify therapeutic targets by highlighting regions, features or specific residues important for T3SS functionality and ii) chimera expression can induce phenotypes (latter linked to cryptic functionality) for further exploration.

### Pathotype-specific T3SS protein functionality

To further support that phenotypes arising from expressing a variant (EPEC, *tarda*, chimera) can be leveraged to describe pathotype-specific functionality, studies were undertaken on a Δ*sepL* mutant phenotype. The latter refers to observing faster migration of proteins through SDS-PA gels (most evident for those >40kDa) with this phenotype rescued by re-introducing EPEC, not *tarda sepL* (Figure 7A). Screening virulence factor-deficient strains revealed the phenotype was shared by non-pathogenic (K12) *E*.*coli* and, surprisingly, another EPEC E2348/69 strain (Yen et al., 2010), hereafter EPEC_J (Figure 7B). Genome sequencing EPEC_J reported a cryptic ∼50Kb deletion (Figure S4) encompassing operons for two virulence-associated exopolysaccharides; O127 antigen and colanic acid. K12 cannot express its O-antigen genes (Blattner et al., 1997) thereby implicating this molecule. The latter was supported by the addition of purified O-antigen rescuing the phenotype (Figure 7B) and anti-O127 western blot signals (Figure 7C). The phenotype is a readout of cellular O-antigen levels by its, direct/indirect, impact on protein migration through SDS-PA gels.

**Figure 7.**
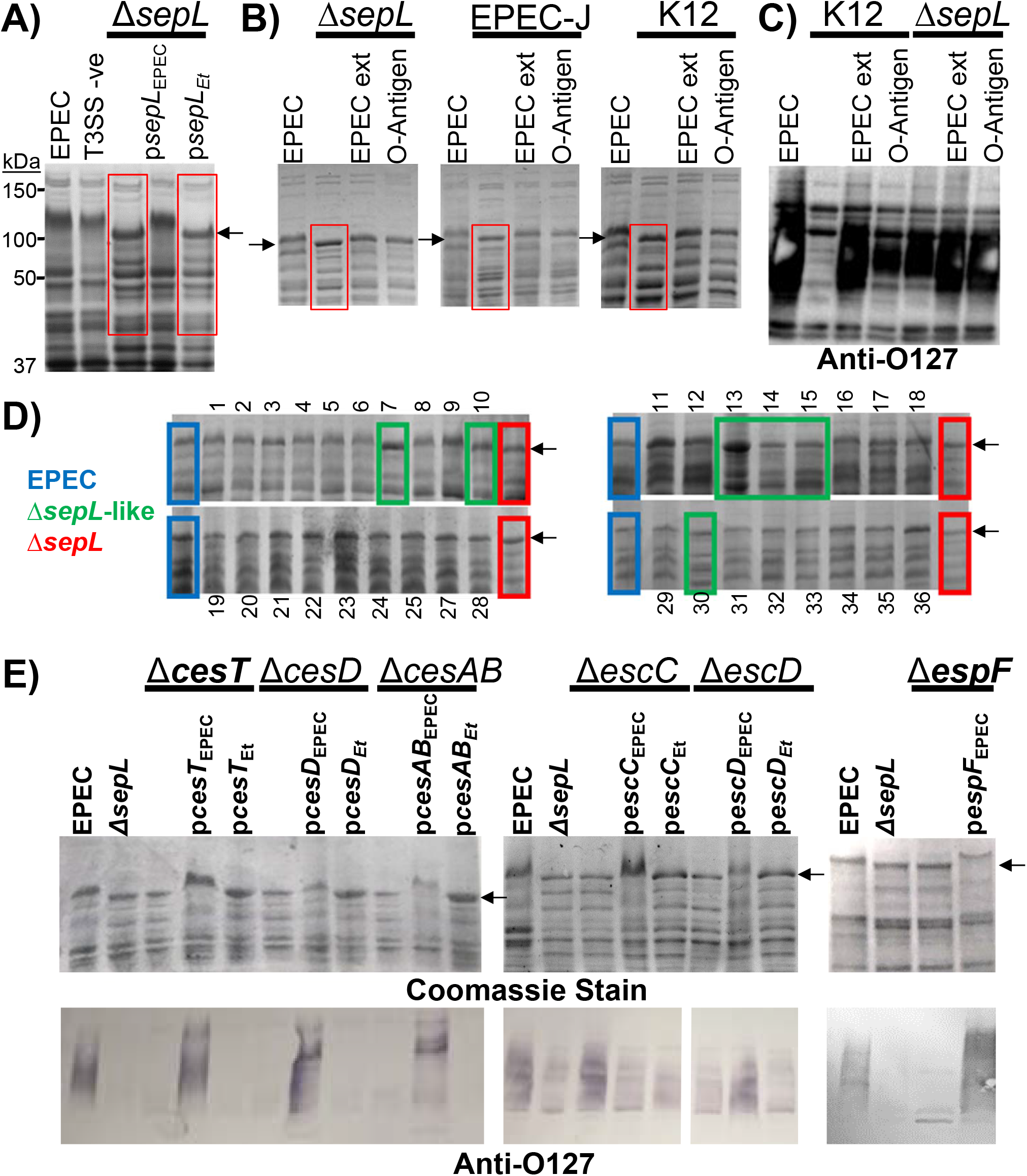
Divergence reflects pathotype-specific functionality. Indicated strains were grown in tissue culture media before isolating total bacterial extracts with samples resolved (SDS-PA gels) for Coomassie blue straining (A, B, D and, top panels, E) or Western blot analysis probing with anti-O127 antibodies (C and, bottom panels, E). EPEC ext. and O-Antigen refers to adding EPEC cellular extracts (1:1 ratio) or commercial O-antigen (1 µg) respectively before analysis. D) Screening EPEC mutants lacking individual LEE genes for *sepL* mutant phenotype. Altered protein migration is highlighted by red rectangles and arrow, with blue and green rectangles highlighting EPEC profile and mutants displaying Δ*sepL* mutant-like phenotype, respectively. Complementation data (E). Strains used were EPEC, EPEC-J (has cryptic 50Kb deletion), non-pathogenic *E*.*coli* (K12) and strains lacking a gatekeeper (Δ*sepL*), a chaperone (Δ*cesT*, Δ*cesD*, Δ*cesAB*), basal body protein (Δ*escD*, Δ*escC*) or effector (Δ*espF*) with or without indicated plasmids (*Et* denotes from *E. tarda*).

Screening a bank of 36 mutants (lack individual LEE genes) reported 17% shared the *sepL* mutant phenotype (Figure 7D). Phenotype rescuing by re-introducing the appropriate EPEC, not *tarda* gene (Figure 7E) demonstrated pathotype-specific functionality, with SepL, for 3 chaperones (CesAB, CesD2, CesT), 2 basal body components (EscD, EscC) and an effector (EspF; no *tarda* homolog). The *tarda* variants fully (CesD, CesAB, EscD, EscC) or partially (CesT, SepL) substituted in T3SS functional assays. These findings provide further evidence for high divergence between LEE homologs reflecting the presence of pathotype-specific biology.

### Other LEE T3SSs

Literature interrogations revealed LEE in other (non-A/E) strains: *Salmonella enterica* subsp. *salamae* (*salamae*) and, a prediction, *Shigella boydii* 13 (Chandry et al., 2012; Walters et al., 2012). Examining the deduced *salamae* protein sequences reveal another, overlooked, example of unprecedented LEE T3SS family divergence; as little as 26.2% and 18.2% identity to EPEC and *tarda* counterparts, respectively (Figure 1B). This finding prompted functionality interchangeability studies, but only for the homologs of the 12 *tarda* complementation-defective genes. These *salamae* proteins are distributed among the 3 homology categories (2, 4 and 6 in lowest, intermediate and highest, respectively; Figure 1B) but each functionally replaced its EPEC counterpart (not shown). This finding emphasizes that LEE proteins can tolerate significant change.

To probe for other LEE-encoding non-A/E strains BLAST searches were undertaken with four EPEC proteins (Tir, EscP, Ler, CesT). This reported Tir and EscP homologs assigned to *Shigella boydii* 13 whose similarity scores (>98%) supported the strain acquiring LEE from EHEC (Walters et al., 2012). The analysis uncovered homologs for all 4 proteins in a recently sequenced strain, *Citrobacter freundii* NCTC6267 (embl accession UFWO01000001.1) with high identity (>82%) to the *salamae* variants indicative of LEE horizontal transfer between these strains.

## Discussion

Uncovering two examples of unprecedented divergence within an evolutionarily closely-related T3SS family has led to the discovery i) that all LEE T3SS proteins can tolerate dramatic change, ii) of tools and strategies to identify therapeutic targets and cryptic, pathotype-specific, functionality, and iv) of novel LEE biology. Moreover, the work provides unique resources and testable predictions with implications for other T3SS families and the identification of anti-virulence (broad- or pathotype-specific) targets.

### LEE *tarda*: Novel biology and functionality

Our interest in *tarda* LEE as a potential research tool prompted sequence data re-interrogation as ∼32% of genes, conserved in A/E pathogens, were absent or truncated (Nakamura et al., 2013). Our analyses revealed *tarda* lacks sequences for 8 (∼20%), not 9, classic LEE genes while four (10%) were not truncated but interrupted (insertion, nonsense or a frameshift mutation). The missing genes are non-essential accessory proteins (Deng et al., 2004; Ruano-Gallego et al., 2015) while each 2-*orf* gene was shown to produce a full length and/or functional protein. Indeed, our finding that all examined *tarda* LEE genes (32 of 33; did not examine non-essential muramidase) produce full length and/or functional proteins (Ler, Tir, EspZ, Intimin data; not shown) provides the first, albeit indirect, evidence for this region encoding a functional injectisome.

The expression/functionality of 2-*orf* genes indicates involvement of ribosome-mediated mechanisms; latter linked to regulating cellular processes including T3SSs (Weiss, 1991). We predict such regulation may functionally replace absence of LEE GrlA/GrlR homologs which modulate EPEC LEE gene transcription in response to environmental signals (Padavannil et al., 2013). The predicted involvement of ribosome-jumping, -readthrough and - slippage mechanisms needs verification, especially in light of finding a 2-*orf* gene (*escD*) did not produce a single polypeptide but independently expressed Orfs. The latter also argues for unusual *tarda* EscD biology as questions how the periplasmic domain (Orf2) could access this compartment and function with Orf1 (cytoplasmic and transmembrane domain) to form a membrane-spanning ring structure (Gaytan et al., 2016). Studies on other *tarda* genes may reveal other non-conventional biology.

### Challenging T3SS dogma

A pivotal discovery was the identification of LEE T3SS proteins from two non-A/E strains (*E*.*tarda* and *S*.*salamae*) displaying unheard levels of divergence relative to their EPEC homolog and in deed each other: as little as 18.2% (*tarda*/*salamae*), 21% (*tarda*/EPEC) and 26.2% (*salamae*/EPEC) identity. The latter contrasts to >80% identity between A/E pathogen homologs (Bhatt et al., 2019; Castillo et al., 2005; Perna et al., 1998). Indeed, the similarity scores (as low as 32.4%) being comparable to those between distinct families (Diepold and Wagner, 2014) suggests *tarda* and *salamae* LEE T3SSs represent distinct subfamilies. A beneficial role for these systems, whether by providing T3SS or other functions, is supported the discovery of closelt-related homologs in other strains: *E*.*tarda* is now a member of new species, *E. anguillarum*, that share, among other things, a conserved LEE region (Nakamura et al., 2013; Shao et al., 2015) with *salamae* LEE homologs in multiple strains (Desai et al., 2013). Illustrating that all 28 examined T3SS proteins (*tarda* or *salamae* variant) can accommodate dramatic change (insertions, deletions, synonymous substitutions) questions the idea that high conservation between A/E pathogen homologs reflects structural constraints (Bhatt et al., 2019; Castillo et al., 2005; Perna et al., 1998). By contrast, the findings argue that evolutionary pressure can in fact dramatically alter each protein with selection of changes most beneficial to the strain’s lifecycle leading to a pathotype-specific T3SS.

### Cryptic T3SS protein biology

The idea that divergence reflects pathotype-specific functionality was first suggested by identifying phenotypes linked to expressing a specific variant (EPEC, *tarda*, chimera). The hypothesis was clearly verified by studies on one phenotype as it revealed key roles for 7 EPEC LEE proteins in controlling the cellular level of another virulence factor, O-127 antigen; latter co-expressed with LEE (Hazen et al., 2015). Crucially, pathotype-specific biology was demonstrated by re-introducing the EPEC gene (not *tarda* variant) rescuing the phenotype; noting the *tarda* variants functionally-substituted (fully or partially) in T3SS assays. These findings not only support the emerging concept of EPEC LEE T3SS proteins possessing cryptic functions (Katsowich et al., 2017; Pal et al., 2019) but extends it by demonstrating that they can possess pathotype-specific biology. We predict that most, if not all LEE T3SS proteins have pathotype-specific functions involved, not in the T3SS export process but to coordinate specific T3SS events with other cellular processes.

Of note, discovering LEE protein/O-antigen crosstalk hinged on unearthing a cryptic undocumented 50Kb deletion in a provided strain (Yen et al., 2010) revealing another example of unintentional genomic alteration (Cepeda-Molero et al., 2017). The latter challenge the EPEC genome stability concept (Nisa et al., 2013) and cautions on the use of these strains and engineered derivatives.

### Therapeutic targets

The discovery that LEE T3SS proteins possess pathotype-specific biology (linked to crosstalk with other cellular processes) opens up avenues of research for uncovering therapeutic targets in T3SS proteins features not critical for T3SS export process or components in processes modulated by the T3SS protein. For example determining how any of the 7 EPEC LEE proteins (a gatekeeper, an effector, 2 basal body components, 3 chaperones) control EPEC O-antigen levels may provide strategies to block such crosstalk. We propose CesT’s role relates to it binding the post-transcriptional regulator, CsrA, which controls many cellular processes including expression of the EPEC T3SS (via binding mRNAs including *sepL*-related) and colanic acid (Berndt et al., 2019; Bhatt et al., 2009; Katsowich et al., 2017). Our work has revealed a myriad of phenotypes that represent future opportunities to uncover additional pathotype-specific biology and/or T3SS crosstalk.

T3SS-specific therapeutic targets were forwarded by domain-swapping experiments (between complementation-defective and EPEC variants) as revealed regions, features or residues important for T3SS functionality. Indeed, such studies can uncover new phenotypes (suggestive of cryptic functionality) and biological insights that may promote therapeutic development. The latter is illustrated studies on the extracellular translocators revealing an EPEC EspA-EspD relationship underpinning a key pathogenic event i.e. stable interaction of the host membrane-inserted Tir effector with EPEC’s Intimin surface protein (Kenny et al., 1997b; Marches et al., 2000). The domain-swapping experiments linked complementation defects to s 10 residue region in EspA - part of an extracellular hypervariable domain (Crepin et al., 2005) (EHVD) - and a specific residue within EspD’s C-terminal coil-coil feature (CCF), not the CCF *per se*. EspD interacts with, and ‘caps’ the EspA filament (Gaytan et al., 2016) with the capping function, based on orthologues (Dey et al., 2019), via EspD’s N-terminal domain. Our findings forward a model whereby EspD transition from its filament ‘capping’ to ‘membrane-inserting’ conformation enables CCF-EHVD interaction needed for efficient Tir delivery and stable Tir-Intimin interaction. Verifying this prediction would provide support this extracellular interaction as a valid therapeutic target. Understanding the key role for Q_354_ within the EspD C-terminal CCF region could promote the development of therapeutic (peptides, small molecule, antibody) reagents. Undoubtedly, similar studies on the remaining 10 complementation-defective *tarda* variants will uncover more phenotypes, pathotype-specific functionality and putative therapeutic targets.

### Research and bioinformatics tools

Uncovering divergent LEE protein homologs that do (16 *tarda* and, undoubtedly, all 28 *salamae* proteins) and do not (12 *tarda* proteins) functionally replace their EPEC counterpart provides a powerful resource for experimental- and bioinformatics-related studies. Probing protein databases for homologs to define ‘natural variation’ can be highly beneficial by highlighting pathotype-specific changes. Divergent homologs can be employed to, for example, i) in co-immunoprecipitation/mass spectrometry studies to reveal partner differences and may also highlight possible crosstalk with another cellular processes while ii) defining *tarda*/*salamae* proteins structures for comparison to those published for A/E homologs (Lyons and Strynadka, 2019; Marshall and Finlay, 2014) may reveal pathotype-specific features as therapeutic targets or for further investigation.

By contrast, comparing the 3 divergent (EPEDC/*tarda*/*salamae*) homologs provides numerous predictions. For example, about a quarter of the T3SS proteins whose homologs differed significantly (10-21 residue) due to terminal extensions or truncations. The latter is suggestive of i) LEE sequence errors, ii) ‘pseudo’ extensions; noting EPEC Ler is expressed from 2^nd^ ATG codon or, more likely, iii) pathotype-specific functionality. Moreover, while homologs for only two proteins (CesT, EscU) had no indels, suggestive of a size-constraint, three (EscP, EscI. EscE) had very large gap, average, scores (20-45%) suggestive of unusual biology. High EscP divergence (Figure S5) at the amino acid (21%-31% identity; 39%-54% similarity) and indel (7%-36% gap; Figure 1B) level, could be useful for interrogating proposed ‘ruler’ and ‘substrate switching’ models; linked to protein size and/or terminal domain features (Gaytan et al., 2016). Another notable aspect was ‘divergence anomalies’ between partners, for example, >74% similarity of SepL homologs versus >42% for CesL chaperone suggesting CesL co-divergence with other protein/s or it has acquired pathotype-specific functionality. Divergence anomalies may also aid understanding on why some T3SS substrates have multiple chaperones, for example, ∼66% similarity of EspD and the CesD2 homologs is suggestive of co-divergence whereas the higher (>77%) value for CesD homologues indicates interacts with more conserved regions of EspD and/or, more likely, other protein/s.

### Constraint-releasing selection pressure?

Why does high divergence in 12 *tarda*, but not corresponding *salamae* proteins compromise T3SS functionality when assessed in EPEC? Our linkage of complementation defects with co-divergence is suggestive of unique pressure for T3SS proteins to tolerate changes which reduce/negate T3SS functionality by co-electing ‘compensatory’ changes in partner protein/s. Possible selection pressures may relate to LEE *tarda*-specific features such as i) 2-*orf* genes, ii) absence of GrlA, GrlR or EspH homologs, or iii) effector repertoire differences: EPEC - 6 LEE and >20 non-LEE-encoded (Nle) effectors (Gaytan et al., 2016); *salamae* - 3 LEE, and possibly a few Nle-like or putative effectors (Desai et al., 2013); *tarda* - 2 LEE but no Nle effectors (Dr Yoji Nakamura; personal communication). Interestingly, adjacent to *tarda* LEE are genes for an effector/cognate chaperone pair (not shown) that are homologous to proteins from another T3SS family (*Yersinia*). We propose this YopH-like effector is a *tarda* LEE T3SS substrate which required dramatic change in the LEE T3SS regulatory proteins to accommodate this effector/chaperone complex and control its timely entry into the LEE T3SS and delivery, via the translocon, into target fish cells.

### Other LEE T3SSs

Blast searches using four EPEC proteins supports the possibility of LEE in other non-A/E strains as the findings are consistent with the prediction of *Shigella boydii* 13 acquiring EHEC LEE (Walters et al., 2012) and suggestive of horizontal transfer of LEE between *S*.*salamae* and, a recently genome sequenced strain, *Citrobacter freundii* NCTC6267. Future sequencing projects may uncover other strains encoding distinct LEE T3SSs to provide additional bioinformatics and experimental tools for promoting the discovery of pathotype-specific functionality, increasing understanding of T3SS protein biology and providing therapeutic (broad- or pathotype-specific) targets.

## Acknowledgements

*E*.*tarda* (strain and LEE DNA sequence) were provided by Dr Yoji Nakamura (Research Center for Fish Diseases, National Research Institute of Aquaculture, Japan Fisheries Research and Education Agency). We thank Dr Wendy Smith and Prof. Anil Wipat (School of Computing Science Newcastle University) for genome sequencing EPEC_J. Research was supported in Kenny lab. through PhD funding (A.M. Ministry of High Education, Libya; B.A. Taibah University, Saudi Arabia) and BG-P lab. with grants from DGAPA, PAPIIT, UNAM (IN212420) and CONACyT (180460).

