## Supplementary File for "Unprecedented Protein Divergence within a T3SS Family"

Methods

Strains, plasmids and primers.

Bacterial strains, plasmids and primers are listed (Tables S1-S3) with strains grown on Luria-Bertani (LB) agar plates or liquid broth containing, when appropriate (Tables S1-S2), antibiotic(s) at final concentrate of 50, 25, 100, 25, 12 and 25 μg/ml for Nalidixic acid (Nal), Kanamycin (Km), Carbenicillin (Cb), Chloramphenicol (Cm), Tetracycline (Tet) and Streptomycin (Step) respectively.

Molecular biology.

EPEC gene knockout mutants were generated using described allelic exchange procedure (Donnenberg and Kaper, 1991). Briefly, appropriate oligonucleotides (Table S3) were used with total EPEC DNA to PCR amplify ~0.5-1Kb upstream and downstream regions of target gene for cloning into a suicide vector, as described (Kenny et al., 1997b; Kenny and Jepson, 2000), for cloning into the suicide vector (Table S2). The suicide vector was used to delete target gene(s) as described (Kenny et al., 1997b; Kenny and Jepson, 2000), with success supported by PCR and loss of T3SS functionality rescued by plasmid re-introducing the appropriate gene. EPEC and *E.tarda* genes were cloned into bacterial expression vectors using gene-specific oligonucleotides (Table S3) and total genomic DNA, with the PCR amplified gene(s) introduced into recipient plasmid via the Gibson Assembly® kit method. Some oligonucleotides included sequences encoding for HA epitope tag or, for domain swap experiments, target gene sequence (Table S3) to generate HA-tagged or *E.tarda*/EPEC chimeric protein-encoding plasmids (Table S2). When appropriate gene or regions thereof were sequenced (Source Bioscience, Cambridge, UK).

Protein secretion assay.

Protein secretion profiles were assessed as previously described (Kenny et al., 1997a; Kenny and Finlay, 1995). Briefly, bacteria were grown in LB (~16h) at 37ºC and diluted (1:10) into high glucose Dulbecco's minimal Eagles medium (DMEM) for growth (6h), 37ºC/5% $CO_2$ atmosphere, before removing bacteria (16,000 x *g*, 5 min) and adding trichloroacetic acid (TCA, 10% final vol.) for 16h at 4ºC. The following day the samples were centrifuged (4°C; 16,000 x *g*, 10 min) with dried pellets resuspended in Laemmli buffer for separation on 12% PA-gels (Laemmli, 1970). Proteins were visualised with Coomassie R-250 blue stain.

Effector translocation assay.

Standard mammalian cell maintenance, infection and fractionation protocols were used as previously described (Kenny et al., 1997b; Kenny and Finlay, 1997). Briefly, HeLa cells (ATCC CCL2) were infected with LB-grown bacteria (~10/host cell) for indicated times before washing (ice cold PBS x2) and adding Triton solution (1% v/v Triton X100 in 50mM PBS [pH 7.5] with, final conc., 0.4 mM $NaVO_4$, 1 mM NaF, and 1x protease cocktail). Centrifugation (16,000 x *g*, 10 min) provided soluble and insoluble (pellet) fractions for separation on SDS-PA gels and Western blot analysis probing with anti-Tir (Kenny et al., 1997b) or, commercial, anti-HA antibodies (Abcam; ab130275).

| Bacterial Strain | Description | Antibiotic | Source |
|---|---|---|---|
| **EPEC E2348/69** | **$Nal^R$ variant** | **$Nal^R$** | **(Levine et al., 1985)** |
| ***Δler*** | **Lacks Ler transcriptional regulator** | **$Km^R$** | **(Bustamante et al., 2011)** |
| ***ΔescE*** | **Lacks EscE (chaperone for EscF)** | **$Nal^R$** | **This study** |
| ***ΔcesAB*** | **Lacks CesAB (chaperone for EspA & EspB)** | **$Strep^R$ /$Km^R$** | **G-P Lab*** |
| ***ΔescK*** | **Lacks sorting platform protein EscK** | **$Strep^R$/$Km^R$** | **(Soto et al., 2017)** |
| ***ΔescL*** | **Lacks sorting platform protein EscL** | **$Strep^R$/$Km^R$** | **(Soto et al., 2017)** |
| ***ΔescRSTU*** | **Lacks export apparatus proteins EscRSTU** | **$Km^R$** | **(Yerushalmi et al., 2014)** |
| ***ΔescU*** | **Lacks export apparatus protein EscU** | **$Strep^R$ /$Km^R$** | **(Soto et al., 2017)** |
| ***ΔcesD*** | **Lacks CesD (chaperone for EspD)** | **$Km^R$** | **(Nadler et al., 2006)** |
| ***ΔescC*** | **Lacks outer membrane ring EscC protein** | **$Nal^R$** | **(Ogino et al., 2006)** |
| ***ΔsepD*** | **Lacks SepD gatekeeper protein** | **$Strep^R$** | **G-P Lab*** |
| ***ΔescJ*** | **Lacks inner membrane ring EscJ protein** | **$Strep^R$/$Km^R$** | **G-P Lab*** |
| ***ΔescI*** | **Lacks T3SS rod (adaptor) EscI protein** | **$Strep^R$/$Km^R$** | **G-P Lab*** |
| ***ΔespZ*** | **Lacks EspZ effector** | **$Nal^R$/$Km^R$** | **This study** |
| ***ΔcesL*** | **Lacks CesL (chaperone for SepL)** | **$Strep^R$/$Km^R$** | **G-P Lab*** |
| ***ΔescV*** | **Lack export apparatus protein EscV** | **$Strep^R$** | **(Gauthier et al., 2003)** |
| ***ΔescN*** | **Lacks EscN ATPase protein** | **$Strep^R$** | **(Gauthier et al., 2003)** |
| ***ΔescO*** | **Lacks EscO ATPase stalk protein** | **$Strep^R$/$Km^R$** | **(Romo-Castillo et al., 2014)** |
| ***ΔescP*** | **Lacks Switch/Ruler EscP protein** | **$Strep^R$/$Km^R$** | **(Monjaras Feria et al., 2012)** |
| ***ΔescQ*** | **Lacks sorting platform protein EscK (C-ring)** | **$Strep^R$ /$Km^R$** | **(Soto et al., 2017)** |
| ***Δtir*** | **Lacks Tir effector** | **$Nal^R$** | **(Kenny et al., 1997b)** |
| ***ΔcesT*** | **Lacks CesT (chaperone of Tir + >10 effectors)** | **$Nal^R$/$Km^R$** | **(Creasey et al., 2003)** |
| ***Δeae*** | **Lacks EPEC surface protein Intimin** | **$Nal^R$** | **(Donnenberg and Kaper, 1991)** |
| ***ΔescD*** | **Lacks inner membrane ring EscD protein** | **$Nal^R$** | **This study** |
| ***ΔsepL*** | **Lacks SepL gatekeeper protein** | **$Strep^R$** | **(Gaytan et al., 2018)** |
| ***ΔespA*** | **Lacks EspA translocator protein** | **$Nal^R$/$Km^R$** | **(Kenny et al., 1996)** |
| ***ΔespD*** | **Lacks EspD translocator protein** | **$Nal^R$/$Km^R$** | **(Lai et al., 1997)** |
| ***ΔespB*** | **Lacks EspB translocator protein** | **$Nal^R$** | **(Taylor et al., 1998)** |
| ***ΔcesD2*** | **Lacks CesD2 (chaperone for EspB/EspD)** | **$Km^R$** | **(Neves et al., 2003)** |
| ***ΔescF*** | **Lacks T3SS needle protein, EscF** | **$Strep^R$/$Km^R$** | **G-P Lab*** |
| ***ΔescG*** | **Lacks EscG (chaperone for EscF)** | **$Nal^R$** | **This study** |
| ***ΔespDB*** | **Lacks EspD and EspB translocators** | **$Nal^R$/$Km^R$** | **This study** |
| ***ΔespAB*** | **Lacks EspA and EspB translocators** | **$Nal^R$/$Km^R$** | **This study** |

**G-P Lab* strains provided from the González-Pedrajo Lab.**

Table S1

| | |
|---|---|
| *ΔescE* FP1 | gcgaccacacccgtcctgtgGTCGTTTGTTAACGAGATGG |
| *ΔescE* RP1 | CTATTCCAGCTCAGTTATCGTTATC |
| *ΔescE* FP2 | CGATAACTGAGCTGGAATAGGAGGATGAGTATTGTGAGCC |
| *ΔescE* RP2 | aaggctctcaagggcatcggCCGAGCTCCTGCCAATAGAATGATAACTGC |
| *escE*$_{Et}$ FP | gcgaccacacccgtcctgtgGCAATCAATCCACAAACTG |
| *escE*$_{Et}$ RP | aaggctctcaagggcatcggGCTCCATTATGTGATCAAATG |
| *escE*$_{EPEC}$ FP | gcgaccacacccgtcctgtgGTGAAGGACACTGAAGAAG |
| *escE*$_{EPEC}$ RP | aaggctctcaagggcatcggCTTGTTTGGCTCACAATACTC |
| *cesAB*$_{Et}$ FP | gcgaccacacccgtcctgtgGATCCTGAACAGTCGACATC |
| *cesAB*$_{Et}$ RP | aaggctctcaagggcatcggGAATTTAGAGATCCACTCAGG |
| *escK*$_{Et}$ FP | gcgaccacacccgtcctgtgCATGGTTATCAACATGGAGAG |
| *escK*$_{Et}$ RP | aaggctctcaagggcatcggGATCTCACTGGCGCTTAAG |
| HA-*escK*$_{Et}$ FP1 | ATGTACCCATACGATGTTCCAGATTACGCTATGAATGGAATAAAAAAAAACGAG |
| HA-*escK*$_{Et}$ FP2 | gcgaccacacccgtcctgtgATGTACCCATACGATGTTCCAG |
| HA-*escK*$_{Et}$ RP | aaggctctcaagggcatcggGATCTCACTGGCGCTTAAG |
| *escL*$_{Et}$ FP | gcgaccacacccgtcctgtgCGTTGCCCAGACTATTTG |
| *escL*$_{Et}$ RP | aaggctctcaagggcatcggGGCTGACTTCCTATAGATATAAG |
| HA-*escL*$_{Et}$ FP1 | ATGTACCCATACGATGTTCCAGATTACGCTATGAAAACAACACGCTACAG |
| HA-*escL*$_{Et}$ FP2 | gcgaccacacccgtcctgtgATGTACCCATACGATGTTCCAG |
| HA-*escL*$_{Et}$ RP | aaggctctcaagggcatcggGGCTGACTTCCTATAGATATAAG |
| *escRSTU*$_{Et}$ FP | gcgaccacacccgtcctgtgGGACATCGATACCATAGTAATGG |
| *escRSTU*$_{Et}$ RP | aaggctctcaagggcatcggGGCTAGTATTTGAGTCCTATC |
| *escU*$_{Et}$ FP | gcgaccacacccgtcctgtgCAGCAGCTAAACGTATTC |
| *escU*$_{Et}$ RP | aaggctctcaagggcatcggGGCTAGTATTTGAGTCCTATC |
| *cesD*$_{Et}$ FP | gcgaccacacccgtcctgtgCAGGAGAAACATAGCGTCATG |
| *cesD*$_{Et}$ RP | aaggctctcaagggcatcgg CAGATGATGAAATTAGCCAG |
| *cesD*$_{EPEC}$ FP | gcgaccacacccgtcctgtgGATAAAATCTGCATCTAGCG |
| *cesD*$_{EPEC}$ RP | aaggctctcaagggcatcgg GATAGAAAATCCCCAGTCAGC |
| *escC*$_{Et}$ FP | gcgaccacacccgtcctgtgGCCGTAGCTATCTAAAGAATG |
| *escC*$_{Et}$ RP | aaggctctcaagggcatcggGCTGTGTCATTGATGTCAATG |
| *escC*$_{EPEC}$ FP | gcgaccacacccgtcctgtgCTAGAGCAATCATCTTCATCAG |
| *escC*$_{EPEC}$ RP | aaggctctcaagggcatcggCCACCATCCTGGTAGAAATC |
| *sepD*$_{Et}$ FP | gcgaccacacccgtcctgtgGTAGCATGCCAATTGTAAAC |
| *sepD*$_{Et}$ RP | aaggctctcaagggcatcgg CGTTGATGAACTCAGCATTG |
| HA-*sepD*$_{Et}$ FP1 | ATGTACCCATACGATGTTCCAGATTACGCTATGAGTAATAGTAATGATTTCTCTATCG |
| HA-*sepD*$_{Et}$ FP2 | gcgaccacacccgtcctgtgATGTACCCATACGATGTTCCAG |
| HA-*sepD*$_{Et}$ RP | aaggctctcaagggcatcggGCTGAGTGTGTTAGCCGATAC |
| HA-*sepD*$_{EPEC}$ FP1 | ATGTACCCATACGATGTTCCAGATTACGCTATGAACAATAATAATGGCATAGCAAAG |
| HA-*sepD*$_{EPEC}$ FP2 | gcgaccacacccgtcctgtgATGTACCCATACGATGTTCCAG |
| HA-*sepD*$_{EPEC}$ RP | aaggctctcaagggcatcggGTTCTTGCCTAATCTTTTTTCC |
| *escJ*$_{Et}$ FP | gcgaccacacccgtcctgtgGCCAGTCTGTTTTCTACATC |
| *escJ*$_{Et}$ RP | aaggctctcaagggcatcggCTGAGAGCTATCACCACTAAC |
| *escI*$_{Et}$ FP | gcgaccacacccgtcctgtgCGGCCAAGCTATAGCTTTG |
| *escI*$_{Et}$ RP | aaggctctcaagggcatcggCTTTGCATCCTGTAATAAGG |
| *cesL*$_{EPEC}$ FP | gcgaccacacccgtcctgtgCCAATACGCTTTTTCAAGC |
| *cesL*$_{EPEC}$ RP | aaggctctcaagggcatcggGCCAGAATAAGATCGTGATATG |
| *cesL*$_{Et}$ FP | gcgaccacacccgtcctgtgCGAGGTCAGTAAGGATGG |
| *cesL*$_{Et}$ RP | aaggctctcaagggcatcggGCCAGAACGAGATCTTGATATTTG |
| *escV*$_{Et}$ FP | gcgaccacacccgtcctgtgGTGAGTTATCACCAGAGCG |
| *escV*$_{Et}$ RP | aaggctctcaagggcatcggCACAACATCATCATGCCTC |
| *escV*$_{EPEC}$ FP | gcgaccacacccgtcctgtgGTACGCAGGGAGTGATTG |
| *escV*$_{EPEC}$ RP | aaggctctcaagggcatcggGCCTTCATATCTGGTAGAC |
| *escV*$_{Et\_EPEC}$ FP1 | GGCATCTATTCCAGGGTTCCCTACATTGGTCTTCTTATTTCTG |
| *escV*$_{Et\_EPEC}$ RP1 | CGAAGTTAGGCTGGTAAGAGgagcactttctgaatacgtgg |
| *escV*$_{Et\_EPEC}$ FP2 | CTCTTACCAGCCTAACTTCG |
| *escV*$_{Et\_EPEC}$ RP2 | GAACCCTGGAATAGATGCC |
| *escN*$_{Et}$ FP | gcgaccacacccgtcctgtgCGTAATATTGACATCGCTCG |
| *escN*$_{Et}$ RP | aaggctctcaagggcatcggGTGTTTCAACATCAGCAAG |
| *escN*$_{Et}$-HA RP1 | CTAAGCGTAATCTGGAACATCGTATGGGTAGCCTACCGTTCTGAATAGTTC |
| *escN*$_{Et}$-HA RP2 | aaggctctcaagggcatcggTTAAGCGTAATCTGGAACATCG |
| *escN*$_{EPEC}$ -HA FP | gcgaccacacccgtcctgtgGATATCAGGCGCTATGTGAAG |
| *escO*$_{EPEC}$ FP | gcgaccacacccgtcctgtgGCACCAAAGATATCAGTAGTTACG |
| *escO*$_{EPEC}$ RP | aaggctctcaagggcatcggCATCTGATGGCCGAAAAG |
| *escO*$_{Et}$ FP | gcgaccacacccgtcctgtgGTCGTCAGCCATTTGATTC |
| *escO*$_{Et}$ RP | aaggctctcaagggcatcggCAGTTGGAGACTCTACTGG |
| *escP*$_{Et}$ FP | gcgaccacacccgtcctgtgTAACTCACTAACGGAAG |
| *escP*$_{Et}$ RP | aaggctctcaagggcatcggCAACCAAATCGATTACAACC |
| *escQ*$_{Et}$ FP | gcgaccacacccgtcctgtgCGCAATAGGATCGAGCATAAG |
| *escQ*$_{Et}$ RP | aaggctctcaagggcatcggCTTTTAGCCGCATAAGCC |
| *escQ*$_{Et}$-HA FP | gcgaccacacccgtcctgtgCGCAATAGGATCGAGCATAAG |
| *escQ*$_{Et}$-HA RP1 | CTAAGCGTAATCTGGAACATCGTATGGGTACTCAAGGTTAAGTAATTCAACATAAAAC |
| *escQ*$_{Et}$-HA RP2 | aaggctctcaagggcatcggTTAAGCGTAATCTGGAACATCG |
| *cesT*$_{Et}$ FP | gcgaccacacccgtcctgtgCGCCAATTAATATCTCGC |
| *cesT*$_{Et}$ RP | aaggctctcaagggcatcggGGAATCAGCTTACTATCTTGG |
| *escD*$_{EPEC}$ FP | gcgaccacacccgtcctgtgCTTAGACGATGTAAGTTCACC |
| *escD*$_{EPEC}$ RP | aaggctctcaagggcatcggCTCCACCCAATAATTTGACATC |
| *escD*$_{Et}$ FP | CGGGATCCGGCTGCATTGATTATTGC BamHI |
| *escD*$_{Et}$ RP | GCGTCGACCCGTTGGTTGTGTCAGG SalI |
| *escD*$_{Et}$ FP | GATGAAGAATCCGGGGCTAC (Used for sequencing *escD* gene) |
| *escD*$_{Et}$ RP | CGTCGCTAGTATCATTACCC (Used for sequencing *escD* gene) |
| *escD*$_{Et}$-HA FP | gcgaccacacccgtcctgtgGGCTGCATTGATTATTGC |
| *escD*$_{Et}$-HA RP1 | CTAAGCGTAATCTGGAACATCGTATGGGTACTTATCATAAGAGATTGGGATATGG |

| | |
|---|---|
| *escD*$_{Et}$-HA RP2 | aaggctctcaagggcatcggTTAAGCGTAATCTGGAACATCG |
| *sepL*$_{Et}$ FR | gcgaccacacccgtcctgtgCTGAGTGCGCTTGTACAG |
| *sepL*$_{Et}$ RP | aaggctctcaagggcatcggCTAGGGTACCGTCTACCTG |
| *sepL*$_{Et}$-HA FP | gcgaccacacccgtcctgtgCTGAGTGCGCTTGTACAG |
| *sepL*$_{Et}$-HA RP1 | CTAAGCGTAATCTGGAACATCGTATGGGTACTCCGAGTCTCTATCTATAACATC |
| *sepL*$_{Et}$-HA RP2 | aaggctctcaagggcatcggTTAAGCGTAATCTGGAACATCG |
| *sepL*$_{EPEC}$-HA FP | gcgaccacacccgtcctgtgGGTGAACTTACATCGTCTAAG |
| *sepL*$_{EPEC}$-HA RP1 | CTAAGCGTAATCTGGAACATCGTATGGGTACATAACATCCTCCTTATAATCTATCAC |
| *sepL*$_{EPEC}$-HA RP2 | aaggctctcaagggcatcggTTAAGCGTAATCTGGAACATCG |
| *espA*$_{Et}$ FP | gcgaccacacccgtcctgtgCAGATGTTATAGATAGAGACTCG |
| *espA*$_{Et}$ RP | aaggctctcaagggcatcggCCAACGGAGGATATAGTTAAATTAG |
| *espD*$_{Et}$ FP | cacacccgtcctgtgGCAATACCGAACGATTTC |
| *espD*$_{Et}$ RP | tctcaagggcatcggGATACCTGACGCGTTATTC |
| *espB*$_{Et}$ FP | CGGGATCCGGCAATGATCTGGTTCG BamHI |
| *espB*$_{Et}$ RP | GCGTCGACCTCATCGCCATCAAGTC salI |
| *espADB*$_{Et}$ FP | aaggctctcaagggcatcggCCAACGGAGGATATAGTTAAATTAG |
| *espADB*$_{Et}$ RP | tctcaagggcatcggCATGGCGAAGATGATATCC |
| *espA*_C1 FP1 | CTCTTACCAGCCTAACTTCG |
| *espA*_C1 RP1 | ATCAATAACGTCATCTTTCGAC |
| *espA*_C1 FP2 | CGAAAGATGACGTTATTGATCTGTTTGAAAAAGTAGGGGC |
| *espA*_C1 RP2 | CGAAGTTAGGCTGGTAAGAGCCAACGGAGGATATAGTTAAATTAG |
| *espA*_C2 FP1 | CTCTTACCAGCCTAACTTCG |
| *espA*_C2 RP1 | CTTCACGGTCTGTAGATCGC |
| *espA*_C2 FP2 | CGATCTACAGACCGTGAAGGCAGCTATTTCAGCTAAAGCG |
| *espA*_C2 RP2 | CGAAGTTAGGCTGGTAAGAGGACCTCACAGACTGGATATCG |
| *espA*_C3 FP1 | CTCTTACCAGCCTAACTTCG |
| *espA*_C3 RP1 | ATCAGCAAACTTTGCAATCG |
| *espA*_C3 FP2 | CGATTGCAAAGTTTGCTGAT*ATGAATGAGGCGTCGAAG* |
| *espA*_C3 RP2 | CGAAGTTAGGCTGGTAAGAGCCAACGGAGGATATAGTTAAATTAG |
| *espA*_C4 FP1 | GCTGCATTAGGGGCAAAG |
| *espA*_C4 RP1 | CTCGGCAAAAACTTTAGTCG |
| *espA*_C4 FP2 | CGACTAAAGTTTTTGCCGAGATGAATGAGGCATCTAAGGAG |
| *espA*_C4 RP2 | CTTTGCCCCTAATGCAGCTTTCACCGTTTGCAAATCAC |
| *espA*_C5 FP1 | ATTGACAATATAAATGCTCAATTAGGCGCTGGCGATCTACAG |
| *espA*_C5 RP1 | TAATTGAGCATTTATATTGTCAATCCCGGAAATTTTTATCTCATTG |
| *espD*_C1 FP1 | CTCTTACCAGCCTAACTTCG |
| *espD*_C1 RP1 | CGGAGCTTTATCACCCATAAC |
| *espD*_C1 FP2 | TTATGGGTGATAAAGCTCCGCAGGCGTTAAAAACAGCAG |
| *espD*_C1 RP2 | cgaagttaggctggtaagagGAAATATCCACTCTGCCATC |
| *espD*_C2 FP1 | CTCTTACCAGCCTAACTTCG |
| *espD*_C2 RP1 | CATATCCAAATTCTTAACGCTAAAC |
| *espD*_C2 FP2 | GTTTAGCGTTAAGAATTTGGATATGCAAAGTGAGTACTTAAGTCAGAGTGC |
| *espD*_C2 RP2 | cgaagttaggctggtaagagGAAATATCCACTCTGCCATC |
| *espD*_C3 FP1 | cacacccgtcctgtgGCAATACCGAACGATTTC |
| *espD*_C3 RP1 | aaggctctcaagggcatcggTTAaactcgaccgctgacaatacggatATTAACACGCGCAGTGTCAG |
| *escF*$_{Et}$ FP | gcgaccacacccgtcctgtgGTACTGAGCAAGAGACTTG |
| *escF*$_{Et}$ RP | aaggctctcaagggcatcggGATTTGCCGTCATACGCTG |
| *escG*$_{EPEC}$ FP | gcgaccacacccgtcctgtgCGTAGAAAGTGGAATGTTGAAAAC |
| *escG*$_{EPEC}$ RP | aaggctctcaagggcatcggGTAGAAGCAGCGTTACTAATTCC |
| *escG*$_{Et}$ FP | gcgaccacacccgtcctgtgGAGAGTGGAATGCTGAAAAC |
| *escG*$_{Et}$ RP | aaggctctcaagggcatcggCAACAGCCCACTACTCTATC |
| *ΔescG* FP1 | gcgaccacacccgtcctgtgCTGACCTTATTAATCGCATG |
| *ΔescG* RP1 | aaggctctcaagggcatcggCTCTTTCTCCACTTTCACCG |
| *ΔescG* FP2 | ataagatactggtgtggtaaGCCAAGATTAGATATAAAGAGGC |
| *ΔescG* RP2 | tctttatatctaatcttggcTTACCACACCAGTATCTTATTAGCAG |
| *ΔescG* FP3 | ggttaaaaaggatcgatcctCTGACCTTATTAATCGCATG |
| *ΔescG* RP3 | gtggaattcccgggagagctCTCTTTCTCCACTTTCACCG |

Primer related features:

**Capital letter** = primer region for amplifying target gene

**Small letter** = 20bp extension for insertion at target region in recipient vector

**Highlighted in Blue** = DNA sequence encoding HA tag

**Highlighted in Red** = DNA sequence for restriction enzyme cleavage site plus extra GC (green)

Table S2

| Plasmid | Description | Antibiotic | Reference |
|---|---|---|---|
| **pCR-2.1** | Cloning vector | Cb$^R$ | Invitrogen |
| **pACYC184** | Cloning vector | Cm$^R$/Tet$^R$ | (Chang and Cohen, 1978) |
| **pTet** | pACYC184 encoding Cm inactivated | Tet$^R$ | Kenny Lab (B Kenny) |
| ***pescE*** | pACYC184 encoding EPEC EscE | Cm$^R$ | This study |
| ***pescE***$_{Et}$ | pACYC184 encoding *E.tarda* EscE | Cm$^R$ | This study |
| ***pcesAB*** | pMQcesAB (pQE30) encoding EPEC His-CesAB chaperone | Cb$^R$ | G-P Lab* |
| ***pcesAB***$_{Et}$ | pACYC184 encoding *E.tarda* CesAB chaperone | Cm$^R$ | This study |
| ***pescK*** | pET encoding EPEC EscK | Cb$^R$ | (Soto et al., 2017) |
| ***pescK***$_{Et}$ | pACYC184 encoding *E.tarda* EscK | Cm$^R$ | This study |
| ***pescL*** | pTrc encoding EPEC EscL | Cb$^R$ | (Soto et al., 2017) |
| ***pescL***$_{Et}$ | pACYC184 encoding *E.tarda* EscL | Cm$^R$ | This study |
| ***pescRSTU***$_{Et}$ | pACYC184 encoding *E.tarda* EscRSTU | Cm$^R$ | This study |
| ***pescU*** | pACYC184 encoding EPEC EscU | Cm$^R$ | This study |
| ***pescU***$_{Et}$ | pACYC184 encoding *E.tarda* EscU | Cm$^R$ | This study |
| ***pcesD*** | pACYC184 encoding EPEC CesD | Cm$^R$ | This study |
| ***pcesD***$_{Et}$ | pACYC184 encoding *E.tarda* CesD | Cm$^R$ | This study |
| ***pescC*** | pACYC184 encoding EPEC EscC | Cm$^R$ | This study |
| ***pescC***$_{Et}$ | pACYC184 encoding *E.tarda* EscC | Cm$^R$ | This study |
| ***psepD*** | pATpD (pTrc99A) encoding EPEC SepD | Cb$^R$ | G-P Lab* |
| ***psepD***$_{Et}$ | pACYC184 encoding *E.tarda* SepD | Cm$^R$ | This study |
| ***pescJ*** | pEEescJ (pET19b) encoding EPEC EscJ | Cb$^R$ | G-P Lab* |
| ***pescJ***$_{Et}$ | pACYC184 encoding *E.tarda* EscJ | Cm$^R$ | This study |
| ***pescI*** | pETeI (pTrc99A) encoding EPEC EscI | Cb$^R$ | G-P Lab* |
| ***pescI***$_{Et}$ | pACYC184 encoding *E.tarda* EscI | Cm$^R$ | This study |
| ***pcesL*** | pACYC184 encoding EPEC | Cm$^R$ | This study |
| ***pcesL***$_{Et}$ | pACYC184 encoding *E.tarda* CesL | Cm$^R$ | This study |
| ***pescV*** | pACYC184 encoding EPEC EscV | Cm$^R$ | This study |
| ***pescV***$_{Et}$ | pACYC184 encoding *E.tarda* EscV | Cm$^R$ | This study |
| ***pescV***$_{Et}$ **chimera** | pACYC184 encoding EscV *E.tarda* (1-344)_EPEC (345-674) | Cm$^R$ | This study |
| ***pescN*** | pAEescN (pET19b) EPEC EscN as His-tagged fusion protein | Cb$^R$ | G-P Lab* |
| ***pescN***$_{Et}$ | pACYC184 encoding *E.tarda* EscN | Cm$^R$ | This study |
| ***pescN*::HA** | pACYC184 encoding EscN as a EscN::HA fusion | Cm$^R$ | This study |
| ***pescN*::HA**$_{Et}$ | pACYC184 encoding EscN *E.tarda* EscN::HA fusion | Cm$^R$ | This study |
| ***pescO*** | pACYC184 encoding EPEC EscO | Cm$^R$ | This study |
| ***pescO***$_{Et}$ | pACYC184 encoding *E.tarda* EscO | Cm$^R$ | This study |
| ***pescP*** | pTrc encoding EPEC EscP | Cb$^R$ | (Monjaras Feria et al., 2012) |
| ***pescP***$_{Et}$ | pACYC184 encoding *E.tarda* EscP | Cm$^R$ | This study |
| ***pescQ*** | pVBescQ (pBAD) EscQ::Myc::His | | G-P Lab* |
| ***pescQ***$_{Et}$ | pACYC184 encoding *E.tarda* EscQ | Cm$^R$ | This study |
| ***pescQ*::**HA$_{Et}$ | pACYC184 encoding *E.tarda* EscQ as EscQ::HA fusion proteins | Cm$^R$ | This study |
| ***pcesT*** | pACYC184 encoding EPEC CesT | Cm$^R$ | Kenny Lab (S Quitard) |
| ***pcesT***$_{Et}$ | pACYC184 encoding *E.tarda* CesT | Cm$^R$ | This study |
| ***pescD*** | pACYC184 encoding EPEC EscD | Cm$^R$ | This study |
| ***pescD***$_{Et}$ | pACYC184 encoding *E.tarda* EscD Orf1 and Orf2 | Cm$^R$ | This study |
| ***pescD orf1_orf2***$_{Et}$**::HA** | pACYC184 encoding EscD Orf1 and Orf2::HA | Cm$^R$ | This study |
| ***psepL*** | pTrc encoding EPEC SepL | Cb$^R$ | (Monjaras Feria et al., 2012) |
| ***psepL***$_{Et}$ | pACYC184 encoding *E.tarda* SepL | Cm$^R$ | This study |
| ***psepL*::HA** | pACYC184 encoding EPEC SepL as SepL::HA fusion protein | Cm$^R$ | This study |
| ***psepL*::HA**$_{Et}$ | pACYC184 encoding *E.tarda* SepL as SepL::HA fusion protein | Cm$^R$ | This study |
| ***pespA*** | pACYC184 encoding EPEC EspA | Cm$^R$ | (Kenny et al., 1996) |
| ***pespA***$_{Et}$ | pACYC184 encoding *E.tarda* EspA | Cm$^R$ | This study |
| ***pespA* C1** | pACYC184 encoding EspA EPEC(1-34)_*tarda*(40-199) | Cm$^R$ | This study |
| ***pespA* C2** | pACYC184 encoding EspA *tarda*(1-145)_EPEC(138-192) | Cm$^R$ | This study |
| ***pespA* C3** | pACYC184 encoding EspA EPEC(1-66)_*tarda*(73-199) | Cm$^R$ | This study |
| ***pespA* C4** | pACYC184 encoding EspA *tarda*(1-70)_EPEC(64-192) | Cm$^R$ | This study |
| ***pespA* C5** | pACYC184 encoding EspA *tarda*(1-128)_EPEC(123-130)_*tarda* (138-199) | Cm$^R$ | This study |
| ***pespD*** | pACYC184 encoding EPEC EspD | Cm$^R$ | Kenny Lab (S Quitard) |
| ***pespD***$_{Et}$ | pACYC184 encoding *E.tarda* EspD | Cm$^R$ | This study |
| ***pespD***$_{Et}$ **C1** | pACYC184 encoding EspD *tarda*(1-230)_EPEC(226-380) | Cm$^R$ | This study |
| ***pespD***$_{Et}$ **C2** | pACYC184 encoding EspD *tarda*(1-336)_EPEC(332-380) | Cm$^R$ | This study |
| ***pespD***$_{Et}$ **C3** | pACYC184 encoding EspD *tarda*(1-377)_EPEC(373-380) | Cm$^R$ | This study |
| ***pespB*** | pACYC184 encoding EPEC EspB | Cm$^R$ | Kenny Lab (S Quitard) |
| ***pespB***$_{Et}$ | pACYC184 encoding *E.tarda* EspB | Cm$^R$ | This study |
| ***pespADB***$_{Et}$ | pACYC184 encoding *E.tarda* EspA, EspB and EspD | Cm$^R$ | This study |
| ***pcesD2*** | pACYC184 encoding EPEC CesD | Cm$^R$ | This study |
| ***pcesD2***$_{Et}$ | pACYC184 encoding *E.tarda* CesD | Cm$^R$ | This study |
| ***pescF*** | pBTeF (pTrc99A) encoding EPEC EscF | Cb$^R$ | G-P Lab* |
| ***pescF***$_{Et}$ | pACYC184 encoding *E.tarda* EscF | Cm$^R$ | This study |
| ***pescG*** | pACYC184 encoding EPEC EscG | Cm$^R$ | This study |
| ***pescG***$_{Et}$ | pACYC184 encoding *E.tarda* EscG | Cm$^R$ | This study |

G-P Lab*: plasmid provided from González-Pedrajo Lab

### Table S3

Bustamante, V.H., Villalba, M.I., Garcia-Angulo, V.A., Vazquez, A., Martinez, L.C., Jimenez, R., and Puente, J.L. (2011). PerC and GrlA independently regulate Ler expression in enteropathogenic Escherichia coli. Mol Microbiol *82*, 398-415.

Chang, A.C., and Cohen, S.N. (1978). Construction and characterization of amplifiable multicopy DNA cloning vehicles derived from the P15A cryptic miniplasmid. Journal of bacteriology *134*, 1141-1156.

Creasey, E.A., Delahay, R.M., Bishop, A.A., Shaw, R.K., Kenny, B., Knutton, S., and Frankel, G. (2003). CesT is a bivalent enteropathogenic *Escherichia coli* chaperone required for translocation of both Tir and Map. Mol Microbiol *47*, 209-221.

Donnenberg, M.S., and Kaper, J.B. (1991). Construction of an *eae* deletion mutant of enteropathogenic *Escherichia coli* by using a positive-selection suicide vector. Infection and immunity *59*, 4310-4317.

Gauthier, A., Puente, J.L., and Finlay, B.B. (2003). Secretin of the enteropathogenic *Escherichia coli* type III secretion system requires components of the type III apparatus for assembly and localization. Infection and immunity *71*, 3310-3319.

Gaytan, M.O., Monjaras Feria, J., Soto, E., Espinosa, N., Benitez, J.M., Georgellis, D., and Gonzalez-Pedrajo, B. (2018). Novel insights into the mechanism of SepL-mediated control of effector secretion in enteropathogenic *Escherichia coli*. Microbiologyopen *7*, e00571.

Kenny, B., Abe, A., Stein, M., and Finlay, B.B. (1997a). Enteropathogenic *Escherichia coli* protein secretion is induced in response to conditions similar to those in the gastrointestinal tract. Infection and immunity *65*, 2606-2612.

Kenny, B., DeVinney, R., Stein, M., Reinscheid, D.J., Frey, E.A., and Finlay, B.B. (1997b). Enteropathogenic *E. coli* (EPEC) transfers its receptor for intimate adherence into mammalian cells. Cell *91*, 511-520.

Kenny, B., and Finlay, B.B. (1995). Protein secretion by enteropathogenic *Escherichia coli* is essential for transducing signals to epithelial cells. Proc Natl Acad Sci USA *92*, 7991-7995.

Kenny, B., and Finlay, B.B. (1997). Intimin-dependent binding of enteropathogenic *Escherichia coli* to host cells triggers novel signaling events, including tyrosine phosphorylation of phospholipase C-gamma1. Infection and immunity *65*, 2528-2536.

Kenny, B., and Jepson, M. (2000). Targeting of an enteropathogenic *Escherichia coli* (EPEC) effector protein to host mitochondria. Cellular microbiology *2*, 579-590.

Kenny, B., Lai, L.C., Finlay, B.B., and Donnenberg, M.S. (1996). EspA, a protein secreted by enteropathogenic *Escherichia coli*, is required to induce signals in epithelial cells. Mol Microbiol *20*, 313-323.

Laemmli, U. (1970). Cleavage of structural proteins during the assembly of the head of bacteriophage T4. Nature *227*, 680-685.

Lai, L.C., Wainwright, L.A., Stone, K.D., and Donnenberg, M.S. (1997). A third secreted protein that is encoded by the enteropathogenic *Escherichia coli* pathogenicity island is required for transduction of signals and for attaching and effacing activities in host cells. Infect Immun *65*, 2211-2217.

Levine, M.M., Nataro, J.P., Karch, H., Baldini, M.M., Kaper, J.B., Black, R.E., Clements, M.L., and O'Brien, A.D. (1985). The diarrheal response of humans to some classic serotypes of enteropathogenic *Escherichia coli* is dependent on a plasmid encoding an enteroadhesiveness factor. The Journal of infectious diseases *152*, 550-559.

Monjaras Feria, J., Garcia-Gomez, E., Espinosa, N., Minamino, T., Namba, K., and Gonzalez-Pedrajo, B. (2012). Role of EscP (Orf16) in injectisome biogenesis and regulation of type III protein secretion in enteropathogenic *Escherichia coli*. Journal of bacteriology *194*, 6029-6045.

Nadler, C., Shifrin, Y., Nov, S., Kobi, S., and Rosenshine, I. (2006). Characterization of enteropathogenic *Escherichia coli* mutants that fail to disrupt host cell spreading and attachment to substratum. Infection and immunity *74*, 839-849.

Neves, B.C., Mundy, R., Petrovska, L., Dougan, G., Knutton, S., and Frankel, G. (2003). CesD2 of enteropathogenic *Escherichia coli* is a second chaperone for the type III secretion translocator protein EspD. Infection and immunity *71*, 2130-2141.

Ogino, T., Ohno, R., Sekiya, K., Kuwae, A., Matsuzawa, T., Nonaka, T., Fukuda, H., Imajoh-Ohmi, S., and Abe, A. (2006). Assembly of the type III secretion apparatus of enteropathogenic *Escherichia coli*. Journal of bacteriology *188*, 2801-2811.

Romo-Castillo, M., Andrade, A., Espinosa, N., Monjaras Feria, J., Soto, E., Diaz-Guerrero, M., and Gonzalez-Pedrajo, B. (2014). EscO, a functional and structural analog of the flagellar FliJ protein, is a positive regulator of EscN ATPase activity of the enteropathogenic Escherichia coli injectisome. Journal of bacteriology *196*, 2227-2241.

Soto, E., Espinosa, N., Diaz-Guerrero, M., Gaytan, M.O., Puente, J.L., and Gonzalez-Pedrajo, B. (2017). Functional Characterization of EscK (Orf4), a Sorting Platform Component of the Enteropathogenic Escherichia coli Injectisome. Journal of bacteriology *199*.

Taylor, K.A., O'Connell, C.B., Luther, P.W., and Donnenberg, M.S. (1998). The EspB protein of enteropathogenic *Escherichia coli* is targeted to the cytoplasm of infected HeLa cells. Infection and immunity *66*, 5501-5507.

Yerushalmi, G., Litvak, Y., Gur-Arie, L., and Rosenshine, I. (2014). Dynamics of expression and maturation of the type III secretion system of enteropathogenic *Escherichia coli*. Journal of bacteriology *196*, 2798-2806.

**i_a)**

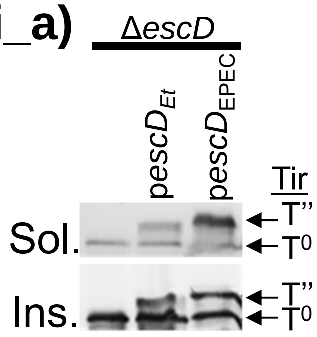


**i_b)** *tarda escD* gene region

```
ATGCTAACCGTACTATCAAGCTACTTTATACTATTAGCAGCAATCATCAACATGCGATCTGATTATAAAATAAGATTTCT
GACTTGGCCACTGAGTGGTAGAGAGTTGATTGTACCTATAGAGTCATTTTCTATTGGTAAAAACGATGATAGTCTCATCA
CATACCAACTTGAAAATCCATTGCAAAATTTCACTCTTAGCACTAATGATGATGGTGTTTGGCTAAATACCCCCACTACA
TTTTGGATTAATGGTATTCAGACGTCTAACGCTGATTCAATTTTGCTTCCACTCAATCAACCAATTGATATTGCCGGCTG
CGGATTTATTCTACTTAGCGCAGATCAAGAAACGTCGCTAGTATCATTACCCAAACGAAAAAAAAACGCACATATAGATA
AGAAAAACATTGTAGTAATCGCATCATTGCTATTTTTAACACTTTCTTTATCGATTTTTGCTTTTTTTATATAAAAAAAC
AAGATTCTATGATTAGTCTCATCAGCAGAGAAGACGTTTATCAATCACTAAAAAAAGAAAAGCTTTTTGCTATTGTACCC
GTCTGGCATGGGAGTAGCATTGCTTTATACGGTCGATGCGAGCAATCATCTCAATTGCAGACTTTCTTTAATTTTCTTTA
TAAAAATAACATAAGATATATAAATAATATTATTTGTAATGACCAAATAATCAGTGGAATTAATGATGTTTTAGTTCAGT
ATGGCTATGACAATATAGATGTATCTTATGGTGGTAGCCCCGGATTCTTCATCCTATCGGGTTACATTCAGTCACCGCTA
CAGTGGAAAAAAGTAGAAGGCCAGCTCTTAACTATGCCAGGCGTCAGAGGATGGCAAGTACGAAACAAAGCCGACGCCAT
CATCAATTCATTAGTTGATGAACTATCGAAAAACAAATTAATCAACAAGTTAAGCATTCATAAGCGTGATAAAGCAATAA
TCATTGACGGCTTAGTTTCAGCAGTAGAAGAGCAAAAAATAATAAATATTATCGACAGGCTAAATAACAATAAAATAATT
TTTCAAAATATACCACCCTATATTCCCAAAAACCTATTTTCAGGAAAGATCATCAGAGTGAGTGGAACAAAAAATGCGCC
AATGATTACATTAGATAATGGGAGCACTCTCACTGTCGGCAGCGCCCTAAACAATGGCTACATAATCAACGAAATCAGCC
TAGAAAATGGTATTAGTATCTCGAGGGAAAATGAATTAATCCATATCCCAATCTCTTATGATAAGTAA
```

### Figure S1

#### ii_a) EscE (35.4%; 51.9%; 12.7%)

```
EPEC        1 MITITELEDEIIKNKEAANVFIEKINDKKNEIHEKMKHPLDKV----TYN      46
              ||:||.|||.||::.|.:.  .|.|...:.|| ||..|.:..:    .|.
E. tarda    1 MISITLLEDIIIEDPEQST--SELIKTLEYEI-EKNNHEMKMIKCNAEYK      47

EPEC       47 EAKELLIACDAAIRTIEIMRIRINNK---      72
              :..:|.||..:||.||..::.||.|:
E. tarda   48 KRDKLNIAYLSAISTITTIKSRIKNEKNR      76
```

## ii_b)

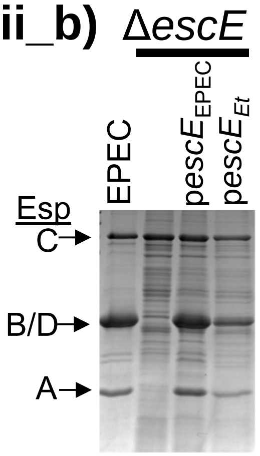


## ii_c)

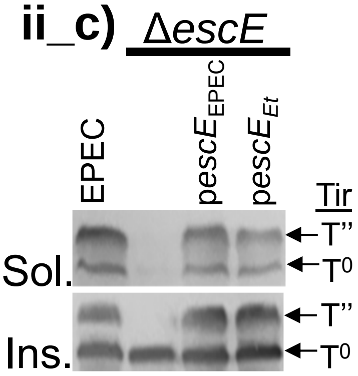


### Figure S1

**iii_a)** **EscG** **(36.8%; 56.8%; 9.5%)**

```
E. tarda    1 MLDGAQRMTANQILIFSAIAAVNHNLRHDAVAMLSALEYVIPNKKDLAQI      50
                 ....::||:||:::|:||.||.|...|.|:|:....:||:|||:|.:
EPEC        1 ---MVNDISANKILVWAAVAAANHKLPKYAEAILNVFPQIIPDKKDIAHL      47

E. tarda   51 ECIILFGLNREDEARQRVSAYADDEISQSLLKICQSGSH------      89
              |.||||||||:::|.:.:....|||.||.|..:......
EPEC       48 EFIILFGLNRKNDAIKALEGCMDDESSQLLYSLFHENVSGWVRGF      92
```

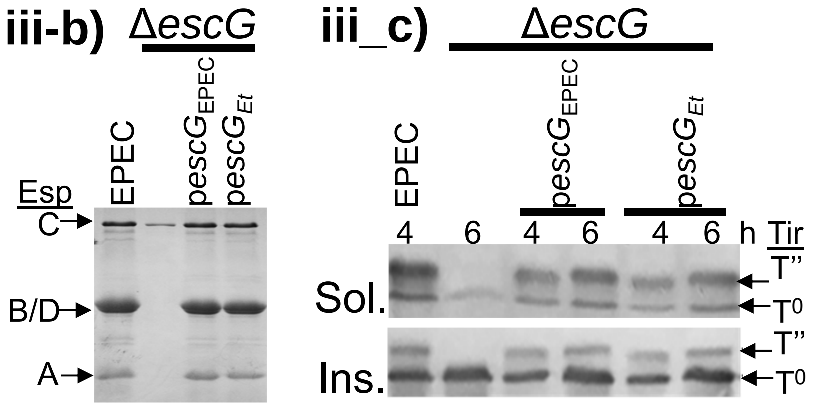


### Figure S1

**iv_a)** **CesD (64.3%; 79.9%; 2.6%)**

```
E. tarda    1 -MNNIKSIQEIIKFYKSGGTLAALTSMTQQDLNVLHGYAYNTFQAGDFTT     49
                .....|:::|..||:.|||||:||::||||||.||.|||..:|:||..|
EPEC        1 MSRKFSSLEDIYDFYQDGGTLASLTNLTQQDLNDLHSYAYTAYQSGDVIT     50

E. tarda   50 ARNIFMLLSYLEHWNYDYTLSLGICHQRLADHEDAQLCFARCATLIMQDP     99
              |||:|.||:||||||||||||||:|||||::||||||||||||||:||||
EPEC       51 ARNLFHLLTYLEHWNYDYTLSLGLCHQRLSNHEDAQLCFARCATLVMQDP    100

E. tarda  100 RAPYYAGISYALTKNNVMAKKAFKACLLWCNGKEEYKSIEDSAKKLLKDC    149
              ||.||:||||.|..|..||||||||||:|||.||:|.:.:::.||||.:.
EPEC      101 RASYYSGISYLLVGNKKMAKKAFKACLMWCNEKEKYTTYKENIKKLLGNT    150

E. tarda  150 KEKT     153
              :
EPEC      151 E---     151
```

**iv_b)**

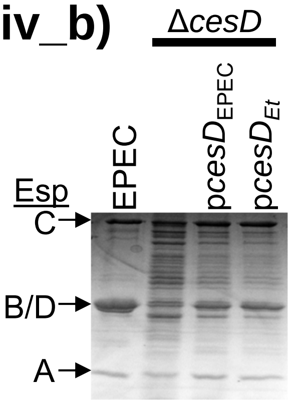


**vi_c)**

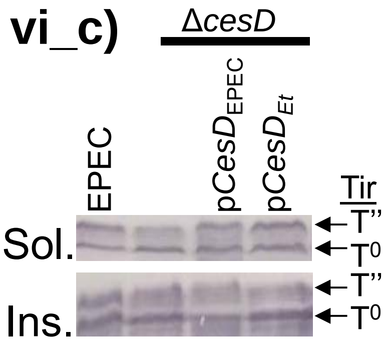


### Figure S1

#### v_a) CesD2 (53.7%; 66.9%; 0.7%)

```
E. tarda     1 MNDTFNDTVINRYLASKGYHLRHEYLMGSGFFIGWCIETSHFSLTYRLDG     50
               |.|||||.|.|.||..|||.::.|:|.||.|||||.|||..|||.||||.
EPEC         1 MVDTFNDEVFNYYLEQKGYTIQKEFLCGSAFFIGWRIETPFFSLAYRLDE     50

E. tarda    51 DELILCYFVSQVNQTGLKSPVLSLSRFLQDLCSRFIEIKVISAMQAMSGN    100
               .|||||.|.:: |||||..|||||:|.|::|...|..||.||||::..|:
EPEC        51 QELILCSFEAR-NQTGLNGPVLSLTRLLEELYHHFSGIKKISAMKSKIGS     99

E. tarda   101 PDEKERRAGLFDFFQSKGARIFSTEQETWFKLFVNN    136
               ..|:::|..||::|..|||....||...||.:.||:
EPEC       100 DSERQKREELFNYFIRKGAVQQETEDGIWFVMNVNS    135
```

**v_b)**

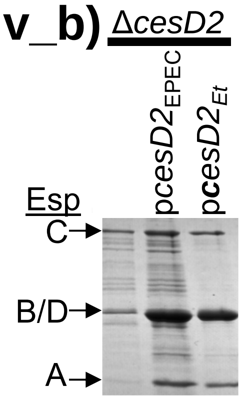


**v_c)**

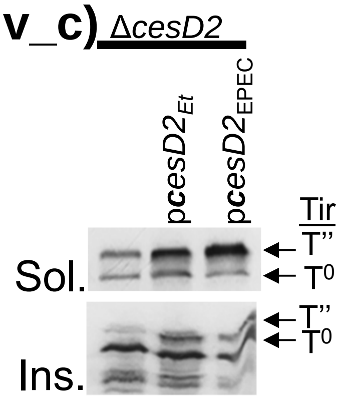


### Figure S1

## vi_a)

**CesL (30.1%; 42.3%; 17.9%)**

```
E. tarda   1 ----MAINENEVCN--NNDVFFV-----VKSNLFDLFIESDADGGVFFTY    39
                 :..|..|...  .||.:..     :..|.|..|||. .|..||.|:
EPEC       1 MNLLVKRNVEEFLRLLGNDFYLFDNRVEIDFNGFSFFIEI-IDNNVFVTF    49

E. tarda  40 SIGYRCDNLVKFLSELSPERINGFIIHVFIYRENLCI-CYQIDDLSSRYE    88
             ::.|..:....|.|.|:|||..|.|.|:|:|...||: |     |.:..:
EPEC      50 ALEYNENAFFSFFSALAPERTQGVIEHIFVYDNKLCLSC-----LLTNID    94

E. tarda  89 KLLLMN-HNKIKSIIERLCR---    107
             ...||| ..:...||||:.|
EPEC      95 VFFLMNTFQQHVQIIERVRRMTS    117
```

#### vi_b) Δ*cesL*

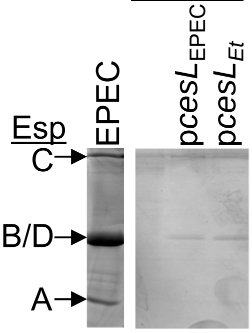


#### vi_c) Δ*cesL*

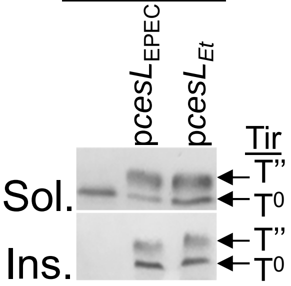


### Figure S1

#### vii_a) **EscR** (73.4%; 88.5%; 0.5%)

```
E. tarda     1 MTQLISIGSQPITLIVIFFLLSLLPIFVVIGTSFLKIVIILGILKNALGI     50
               |:||::||||||.||::||||||||||||||||||||.|:||||||||||
EPEC         1 MSQLMTIGSQPIFLIIVFFLLSLLPIFVVIGTSFLKISIVLGILKNALGI     50

E. tarda    51 QQVPPNMAITSVALTLTIFIMSPTILQVKKNLETNPIDINNNNYFESVDK    100
               ||||||||:|||:|.||:|||||.|||:..|:...||:..::::|:.||:
EPEC        51 QQVPPNMALTSVSLILTMFIMSPIILQINDNISQEPINYTDSDFFQKVDE    100

E. tarda   101 NILSPYKDFLIKNTDVKKIDFFEKKAQEKWGAHSDVASKESIFILLPAFT    150
               .|||||:.||.|||:...::|||:.||:|.| :..:..|:|:||||||||
EPEC       101 KILSPYRGFLEKNTEKDNVEFFERAAQKKLG-NETILKKDSLFILLPAFT    149

E. tarda   151 LGQLESAFKIGFLLYLPFIAIDLIISNILLALGMMMVSPITISIPFKILL    200
               :||||:|||||||||||||||||||||||||||||||||:||||||||||
EPEC       150 MGQLEAAFKIGFLLYLPFIAIDLIISNILLALGMMMVSPVTISIPFKILL    199

E. tarda   201 FILIGGWQKLFEYLLVVN    218
               |||:||||||||:|||||
EPEC       200 FILVGGWQKLFEFLLVVN    217
```

#### vii_b) Δ*escRSTU*

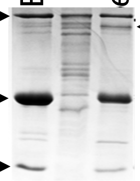


#### vii_c) Δ*escRSTU*

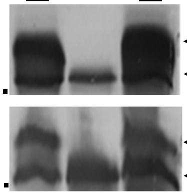


### Figure S1

#### viii_a) EscS (63%; 82.6%; 6.5%)

```
E. tarda  1 MNSGTLVQVSIEMLWVVLLLSLPTVAAASIVGILISLIQALTQLQDQTLP     50
            |::|..||:.::..|::.:||||||.|||::||:|||:||:|||||||||
EPEC      1 MDTGYFVQLCVQTFWIIFILSLPTVIAASVIGIIISLVQAITQLQDQTLP     50

E. tarda 51 FLLKILAVFATLALTYHWMGNVILNFSSLIFEHLNLLVK---     89
            |||||:||||||||||||||..|:||||:|||   ::.|
EPEC     51 FLLKIIAVFATLALTYHWMGTTIINFSSIIFE---MIPKVNG     89
```

#### viii_b) Δ*escRSTU*

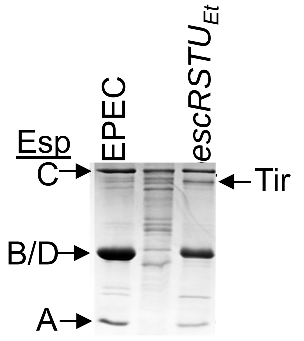


#### viii_c) Δ*escRSTU*

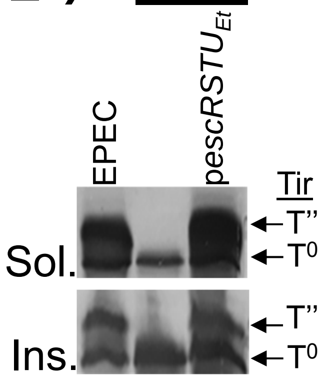


### Figure S1

**ix_a)** **EscT (56.1%; 75.6%; 2.7%)**

```
E. tarda    1 MMKDTVDIIITIFICILRPLGAFLIIPIFSLGNFLSPLIRNAFTLAISLP     50
               |.:.:.:|::.|.|||||||.|:|:||||.|..||..|||:..:|.:||
EPEC        1 -MNEIMTVIVSSFYCILRPLGMFIILPIFSTGVLLSNFIRNSIMIAFTLP     49

E. tarda   51 IIHQNINITEVMPHEISSLSLFVLKELMIGFFIGLSFTIIFWAIDSASQL    100
              ||.:|...:|.:|..|..|:..||||:.|||||||||||:|||||:|.|:
EPEC       50 IIVENYTFSEKLPSGIFQLTGIVLKEISIGFFIGLSFTILFWAIDAAGQI     99

E. tarda  101 IDTLRGSTIASILNPAINDSSSVTGVFLYHFVNIIFIIHGGIEDILSTLY    150
              |||||||||:||.||:|:||||:|||.||.|:::||:||||||:.||..||
EPEC      100 IDTLRGSTISSIFNPSISDSSSITGVILYQFISVIFVIHGGIQSILDKLY    149

E. tarda  151 SSYQTLPISQSIEINGQLISFIYSLWNSFLKLLVSFSIPMIVGILLTDIA    200
              .||:.||:...|..|..||.|::|||:||:||::|||:|||:||.|.|:.
EPEC      150 LSYEILPLQADIAFNRALIDFLFSLWDSFIKLMLSFSVPMIIGIFLCDMG    199

E. tarda  201 FGFLNKTAQQLNVFTLSLPIKSLIAFFILIIVLHTYPSLVKEAI---ALN    247
              ||||||||.||||||||||:||||||.|||::|:|.:|..:...|   .::
EPEC      200 FGFLNKTAPQLNVFTLSLPVKSLIAIFILLLVIHVFPDFITANIHSDIID    249

E. tarda  248 KNLITLLLEKAI    259
              ::|.:::.|
EPEC      250 RSLPSMINE---    258
```

**ix_b)** Δ*escRSTU* **ix_c)** Δ*escRSTU* **ix_d)** Δ*escU*

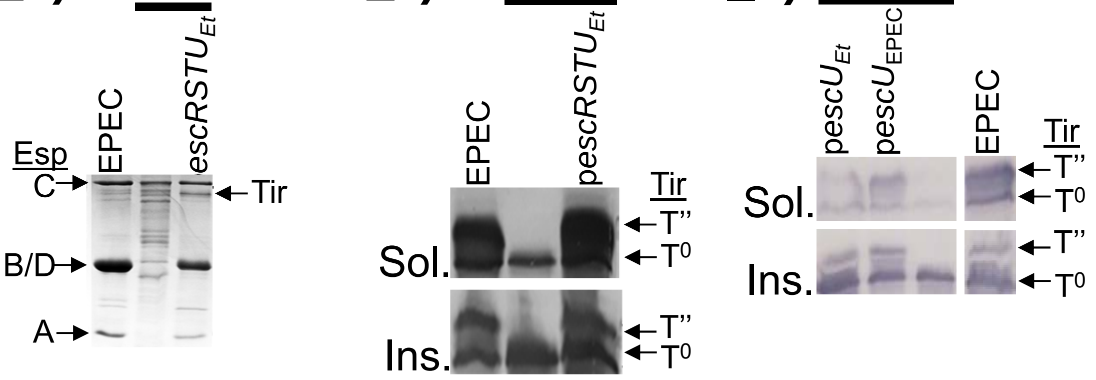


### Figure S1

**x_a)** **EscU** **(58%; 78.8%; 0%)**

```
E. tarda    1 MSEKTEKPTTKKLRDLKEKGDVIKSEEVVSAVQSIFIFTYLYLYGNSFLS     50
              |||||||||.|||||||:||||.|||||::||||:.:|::..|||.||..
EPEC        1 MSEKTEKPTPKKLRDLKKKGDVTKSEEVMAAVQSLILFSFFSLYGMSFFV     50

E. tarda   51 EIIELINTSIESINYELSYSAGKITGMALDLSIKYILPLVAVIFIGDILS    100
              :|:.|:||:|:|:|....|:..:|.|..|::.:.||||:..::|:|.:.:
EPEC       51 DIVGLVNTTIDSLNRPFLYAIREILGAVLNIFLLYILPISLIVFVGTVTT    100

E. tarda  101 IVSQIGFVFAVEKIKPSLQKLSVKNNIKNIFSLKNVFELLKSILKLAFIS    150
              .||||||:|||||||||.||:|||||:|||||:|::||||||:.||..|.
EPEC      101 GVSQIGFIFAVEKIKPSAQKISVKNNLKNIFSVKSIFELLKSVFKLVIIV    150

E. tarda  151 LVSYVIIREHVRDFSNLPYASNTVAFDYSFYIISLLWKGILVGYLIFSIF    200
              |:.|.:...:..:|:|....:...|.....:.:.|||||:|.|||:||:|
EPEC      151 LIFYFMGHSYANEFANFTGLNAYQALVVVAFFVFLLWKGVLFGYLLFSVF    200

E. tarda  201 DFWFQRRNGEKKIMMTKDEVKRESKDSDGNPEVKSERKKIHAEIQSGSLG    250
              |||||:..|.||:.|:|||||||:||:|||||:|.||:::|:|||||||.
EPEC      201 DFWFQKHEGLKKMKMSKDEVKREAKDTDGNPEIKGERRRLHSEIQSGSLA    250

E. tarda  251 NNVRKSTVIVKNPTHIAVCIYYKLGETPVPQILEIGIGKLALHILKIAED    300
              ||::|||||||||||||:|:||||||||:|.::|.|....||.|:|:||.
EPEC      251 NNIKKSTVIVKNPTHIAICLYYKLGETPLPLVIETGKDAKALQIIKLAEL    300

E. tarda  301 NNIPIVEEISLARGIHANIAPGDYITEDFFEPVAEILRIAMNLDY     345
              .:||::|:|.|||.::.||..|.|||||||||||:::|||::|||
EPEC      301 YDIPVIEDIPLARSLYKNIHKGQYITEDFFEPVAQLIRIAIDLDY     345
```

**x_b)** Δ*escRSTU*

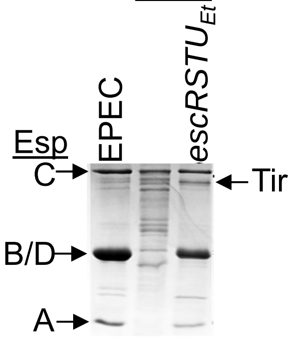


**x_c)** Δ*escRSTU*

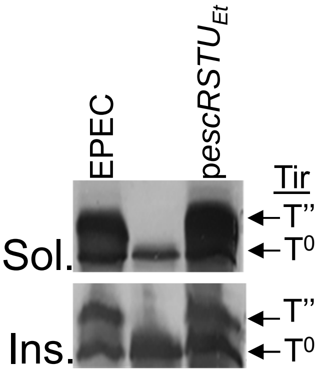


**x_d)** Δ*escU*

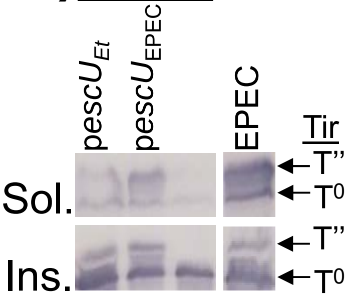


### Figure S1

#### xi_a) EscJ (53.1%; 72.9%; 7.8%)

```
E. tarda     1 ----------MIFIFLITGCKEILYSELSEAEANQMQALLLTNNIDVDKT     40
                         :..|.|...|||.||:.|:|.||||||||||:|:::|.|.
EPEC         1 MKKHIKNLFLLAAICLTVACKEQLYTGLTEKEANQMQALLLSNDVNVSKE     50

E. tarda    41 IEKSKAISLSVDKKDFVKSIKILNDNGFPKKKLVSIETIFPASQLISSPM     90
               ::||..::|||:|:|||::|.|||:|||||||...||.|||.|||::||.
EPEC        51 MDKSGNMTLSVEKEDFVRAITILNNNGFPKKKFADIEVIFPPSQLVASPS    100

E. tarda    91 QESAKLNFIKEQSIEKFLNKIPGVVDSSVTLNALKTDGSSGVSTASVLII    140
               ||:||:|::|||.||:.|:|||||:|.||:||.  .:..|..|:|:||:|
EPEC       101 QENAKINYLKEQDIERLLSKIPGVIDCSVSLNV--NNNESQPSSAAVLVI    148

E. tarda   141 TAPQANLEPSINQIKGLVKNSVDELSIENITVVIKKLAI---    179
               ::|:.||.||:.|||.|||||||:|.:|||:||||..:.
EPEC       149 SSPEVNLAPSVIQIKNLVKNSVDDLKLENISVVIKSSSGQDG    190
```

## xi_b)

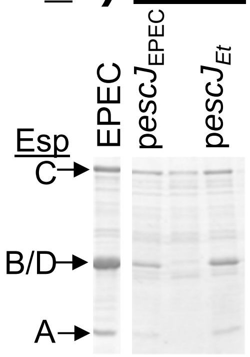


## xi_c)

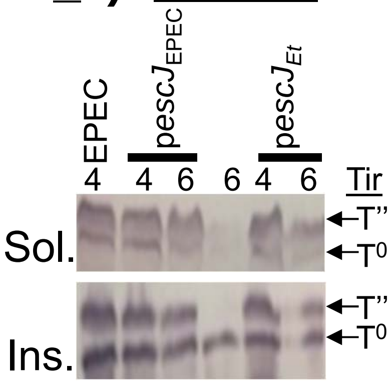

#### xii_a) **EscI** (46.9%; 63.3%; 15%)

```
E. tarda   1 -----------------MDPILQTNALSSILPHSAEASTIPSSVQLTNMP     33
                              |:.|.|:..::..|..:::.:.:...:.|.:.|
EPEC       1 MDALCYCLSHEKRLTVNMNNINQSENINIQLNKASQTNVVDEHIPLASTP     50

E. tarda  34 AQPVEMTAASQFLSQLLPSTASVASPEQVLIEEIKRRHFNALNAPDLNFD     83
             :    ...|:|||.||||.||.|:|||||||||||:||...:|. ||:||
EPEC      51 S----AAGAAQFLDQLLPKTAGVSSPEQVLIEEIKKRHLATMNT-DLSFD     95

E. tarda  84 DLAAGKLSPTDMLQLQRSVLNANVNIDVISKLASLFSTSITKLTSMQ    130
             .|:||.|||.|:|.||::|||||||:||:||||||.|||:|||.|||
EPEC      96 ALSAGGLSPEDVLTLQKNVLNANVNVDVVSKLASLLSTSVTKLVSMQ    142
```

#### xii_b) Δ*escI*

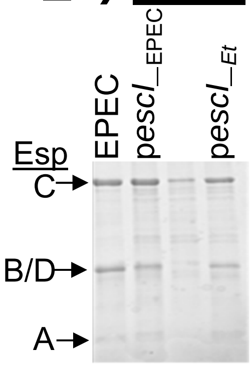


#### xii_c) Δ*escI*

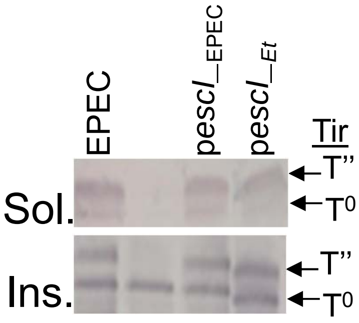


### Figure S1

#### xiii_a) EscC (64.6%; 80.3%; 1.8%)

```
E. tarda     1 MKKSYIFLFTIGMLPYLNVSANTLSQRLA-GDYFIVTPKTDVSIIINDIA    49
               |||...|:||.........:.::|.:||. .:|||:|..:.|..|:||.|
EPEC         1 MKKISFFIFTALFCCSAQAAPSSLEKRLGKNEYFIITKSSPVRAILNDFA    50

E. tarda    50 ANYSLPVFISPKITDSFLGELKADNALIILDKLAKMYHLSLYYDENILYV    99
               ||||:|||||..:.|.|.||:|.:..:.:|:||:|:|||:.||||||||:
EPEC        51 ANYSIPVFISSSVNDDFSGEIKNEKPVKVLEKLSKLYHLTWYYDENILYI   100

E. tarda   100 YKSDEITRSIITPNYLDTNTLISYLKKPTQNNNASCQIKKISSFNSLEVR   149
               ||::||:||||||.|||.::|:.||......|..||.::||::|||:|||
EPEC       101 YKTNEISRSIITPTYLDIDSLLKYLSDTISVNKNSCNVRKITTFNSIEVR   150

E. tarda   150 GVPECIKYITNLASNLDKESQTKATNKDVIRVFKLKYASAVDINYKYRDQ   199
               ||||||||||:|:.:||||:|:||.||||::||||.||||.||.||||||
EPEC       151 GVPECIKYITSLSESLDKEAQSKAKNKDVVKVFKLNYASATDITYKYRDQ   200

E. tarda   200 SVTVPGVISILKTMANDGTLPSQGNNATDSSKLLSGGQAQISADPRQNAI   249
               :|.||||:||||||||::|:|||.|..|.:.|..|......||||||.||:
EPEC       201 NVVVPGVVSILKTMASNGSLPSTGKGAVERSGNLFDNSVTISADPRLNAV   250

E. tarda   250 IVKDREITMDIYQQLITHLDVEQRQIEISVSIIDVDANDLQQLGVNWSGS   299
               :|||||||||||||||:.||:||||||||||||||||||||||||||||:
EPEC       251 VVKDREITMDIYQQLISELDIEQRQIEISVSIIDVDANDLQQLGVNWSGT   300

E. tarda   300 MSFGNSSVSFNSQNTSATLSTAVISNANNFLIRVNALQKNSKAKILSQPS   349
               ::.|..:::|||....|.:|::|||||:||:|||||||:|||||||||||
EPEC       301 LNAGQGTIAFNSSTAQANISSSVISNASNFMIRVNALQQNSKAKILSQPS   350

E. tarda   350 VITLNNMQAILDKNVTFFTKVSSERSASLESITSGTLLRVTPRIINSQNN   399
               :||||||||||||||||:||||.|:.||||||||||||||||||::..:|
EPEC       351 IITLNNMQAILDKNVTFYTKVSGEKVASLESITSGTLLRVTPRILDDSSN   400

E. tarda   400 SDVLK-NEKVRLLLDIQDGSQSPSTQNNQEA---LPQVQNSEITTEATLN   445
               |...| .|:|||||||||||:||.:..|.|:|   ||:||||||:||||||:
EPEC       401 SLTGKRRERVRLLLDIQDGNQSTNQSNAQDASSTLPEVQNSEMTTEATLS   450

E. tarda   446 AGESLLLGGFIQDREISGKAGIPLLRDIPLIGHLFSSSTQEKHSVMRLFL   495
               |||||||||||||:|.|.|.|||||.|||:||.||||:.::||||:||||
EPEC       451 AGESLLLGGFIQDKESSSKDGIPLLSDIPVIGSLFSSTVKQKHSVVRLFL   500

E. tarda   496 IKATPINN----    503
               ||||||.:
EPEC       501 IKATPIKSASSE    512
```

#### xiii_b) ΔescC

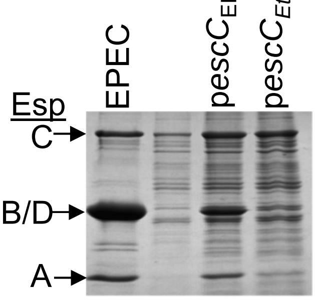


#### xiii_c) ΔescC

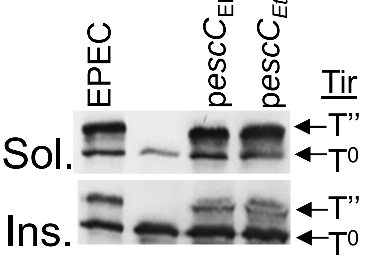


### Figure S1

#### xiv_a) EscF (67.5%; 87.8%; 1.4%)

```
E. tarda  1 MNLIDVTTQMGQFGQRISDSIQPLLASGEDVVNNPERMLQLQFAVQQYSA  50
            |||.::|.|||:.|:.:|||: |.|.:..|:||:||:||:||||||||||
EPEC      1 MNLSEITQQMGEVGKTLSDSV-PELLNSTDLVNDPEKMLELQFAVQQYSA  49

E. tarda 51 YVNVESGMLKTVKDIISTIANRSF  74
            |||||||||||:||::|||:||||
EPEC     50 YVNVESGMLKTIKDLVSTISNRSF  73
```

xiv_b) ΔescF xiv_c) ΔescF

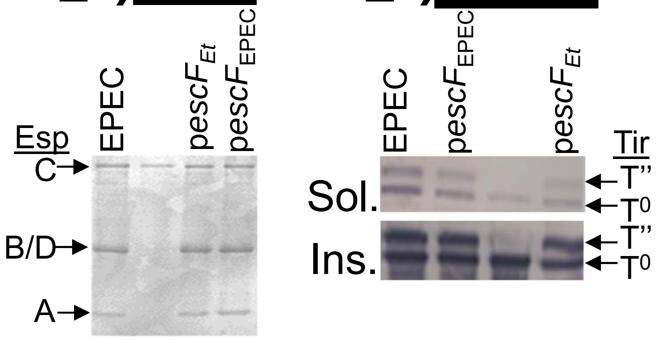


### Figure S1

**xv_a)** **EscO (25%; 54.5%; 12.9%)**

```
EPEC        1 MLDRILSIRKSRANRLRESMAKINSQIKEVDGKLDDCEQSIKESIASKQA
              ||:.||:|:|.|..:.:::|.:.:|.::....||..::.:..:|...:.
E. tarda    1 MLNDILAIKKRRILKKKKNLADVETQKQQAFIDLDTYQRRLTSNIQVYKN

EPEC       51 YCASLVNLDKVSLYKYQIKNNAFD-EQKQRLYEKKSS------LSKEKRS
              :|.:|.::::.:||::|:.|...|: :.||.:.:||..      |||...|
E. tarda   51 FCDNLTSIEFISLFEYRKKQADFEYDMKQLILDKKECENNICVLSKNINS

EPEC       94 LLDSQKRTKENLQHVNKSVEKLSFAIKEHYFD    125
              |       .|:::.:|.|:||:.:.:.:.
E. tarda  101 L-------TEDIKKINISIEKIKYVLNDE---    122
```

**xv_b)**

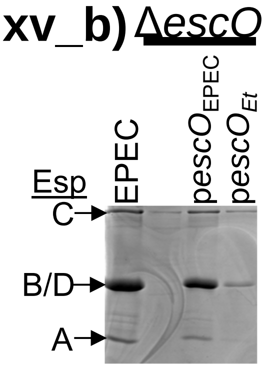


**xv_c)**

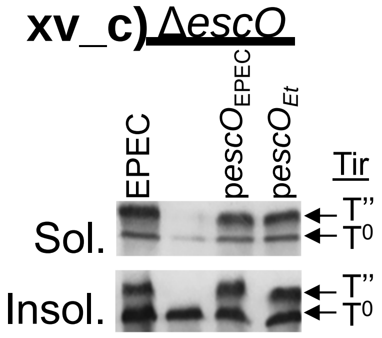


### Figure S1

**i_a)** **EspB** **(37.8%; 55.7%; 12.2%)**

```
EPEC        1 -MNTIDNN-NAAIAVNSVLSSTTDSTSSTTTSTSSISSSLLTDGRVDISK   48
               :||:|.: |.|..:|||         :|....|...|..|..|.:|:::
E. tarda    1 MINTVDGSANNASGINSV---------NTALGVSPTPSETLPAGALDLNE   41

EPEC       49 LLLEVQKLLREMVTTLQDYLQKQLAQSYDIQKAVFESQNKAIDEKKAGAT   98
              |||::|.|||:.:..||:|.|:|:.||:.||:|.|.||:|||:|::.|||
E. tarda   42 LLLQIQVLLRKALRVLQEYQQQQVGQSFKIQEAAFASQDKAIEERRKGAT   91

EPEC       99 AALIGGAISSVLGILGSFAAINSATKGASDVAQQAASTSAKSIGTVSEAS  148
              .|||||.|.|.||:|||||.|..|.:...  |..||:..|.|.|.|:...
E. tarda   92 TALIGGIIGSTLGVLGSFAGITQAREAVK--AGSAAAKIADSTGDVTSEL  139

EPEC      149 TKALAKASEGIADAADDAAGAMQQTIATAAKAASRTSGITDDVATSAQ--  196
              .|:.|:.|: :||||.|.|.:.||.::..|...||.|.|.||:|.:.|
E. tarda  140 AKSTAQLSK-VADAAADTAQSTQQALSRTASITSRASDIADDMAQNTQQA  188

EPEC      197 ---------KASQVAEEAADAAQELAQKAG-----------LLSRFTAAA  226
                       :|:.|||:|..:...:|..|.           |.||....|
E. tarda  189 VSRAASLTRRAADVAEDAVTSTAPMAAVANDVASATDDVVELSSRLKCMA  238

EPEC      227 -------GRISGSTPFIVVTSLAEGTKTLPTTISESVKSNHDINEQRAKS  269
                     ..||.:..||....||:|.|.||.|||..:|.::|:...|||:
E. tarda  239 ESVTNKFDSISQNGAFIAGVHLAQGVKELPGTISAGLKVSNDLAADRAKN  288

EPEC      270 VENLQASNLDTYKQDVRRAQDDISSRLRDMTTTARDLTDLINRMGQAARL  319
              :|:.|..:.:.|:|||:.::|::..||.|:|...|::.|::.|.|||.|:
E. tarda  289 LEDYQQQSRNIYQQDVQGSKDEVRQRLNDITEVTRNINDILTRQGQAVRI  338

EPEC      320 AG    321
              ||
          339 AG    340
```

### Figure S2c

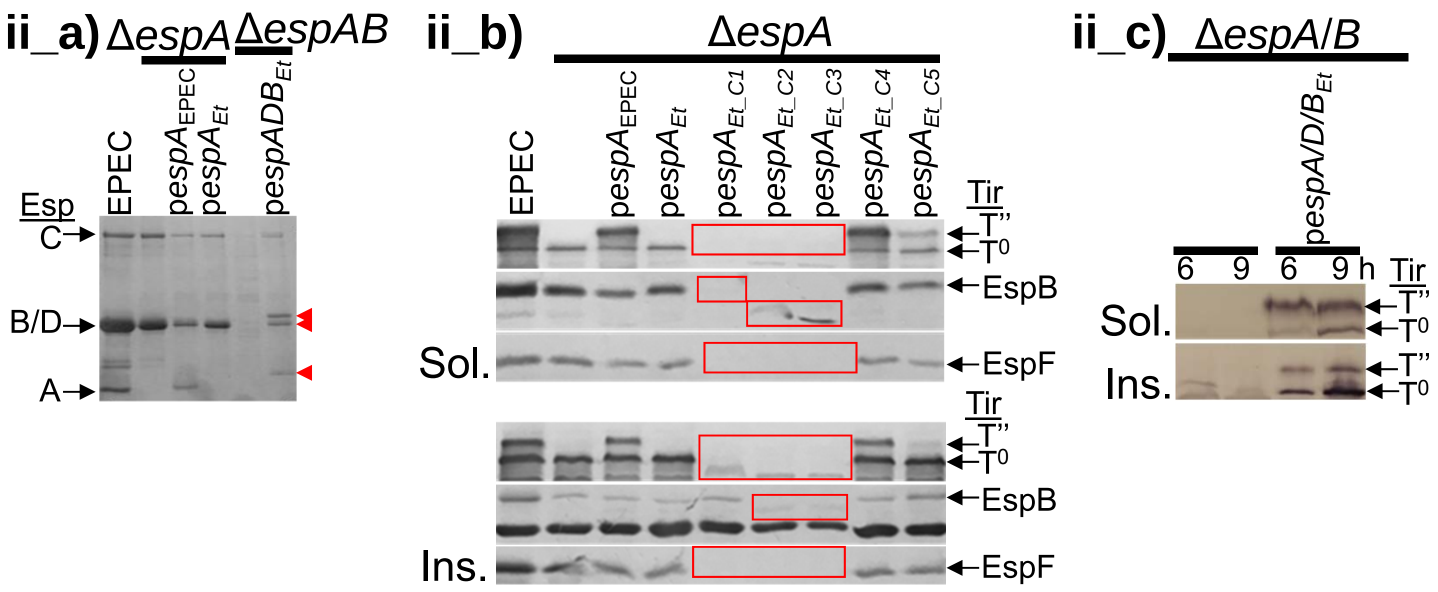
ii_a) ΔespA ΔespAB
EPEC
pespA_EPEC
pespA_Et
pespADB_Et
Esp
C
B/D
A
ii_b) ΔespA
EPEC
pespA_EPEC
pespA_Et
pespA_Et_C1
pespA_Et_C2
pespA_Et_C3
pespA_Et_C4
pespA_Et_C5
Tir
T''
T0
EspB
EspF
Sol.
Ins.
ii_c) ΔespA/B
pespA/D/B_Et
6 9 6 9 h
Sol.
Ins.

### Figure S2

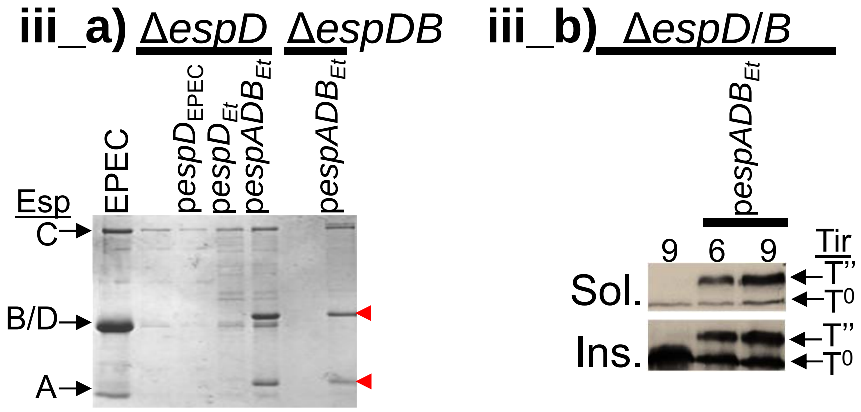
iii_a)
ΔespD
ΔespDB
Esp
EPEC
pespD_EPEC
pespD_Et
pespADB_Et
pespADB_Et
C
B/D
A
iii_b)
ΔespD/B
pespADB_Et
9
6
9
Tir
Sol.
Ins.
T''
T0
T''
T0

### Figure S2

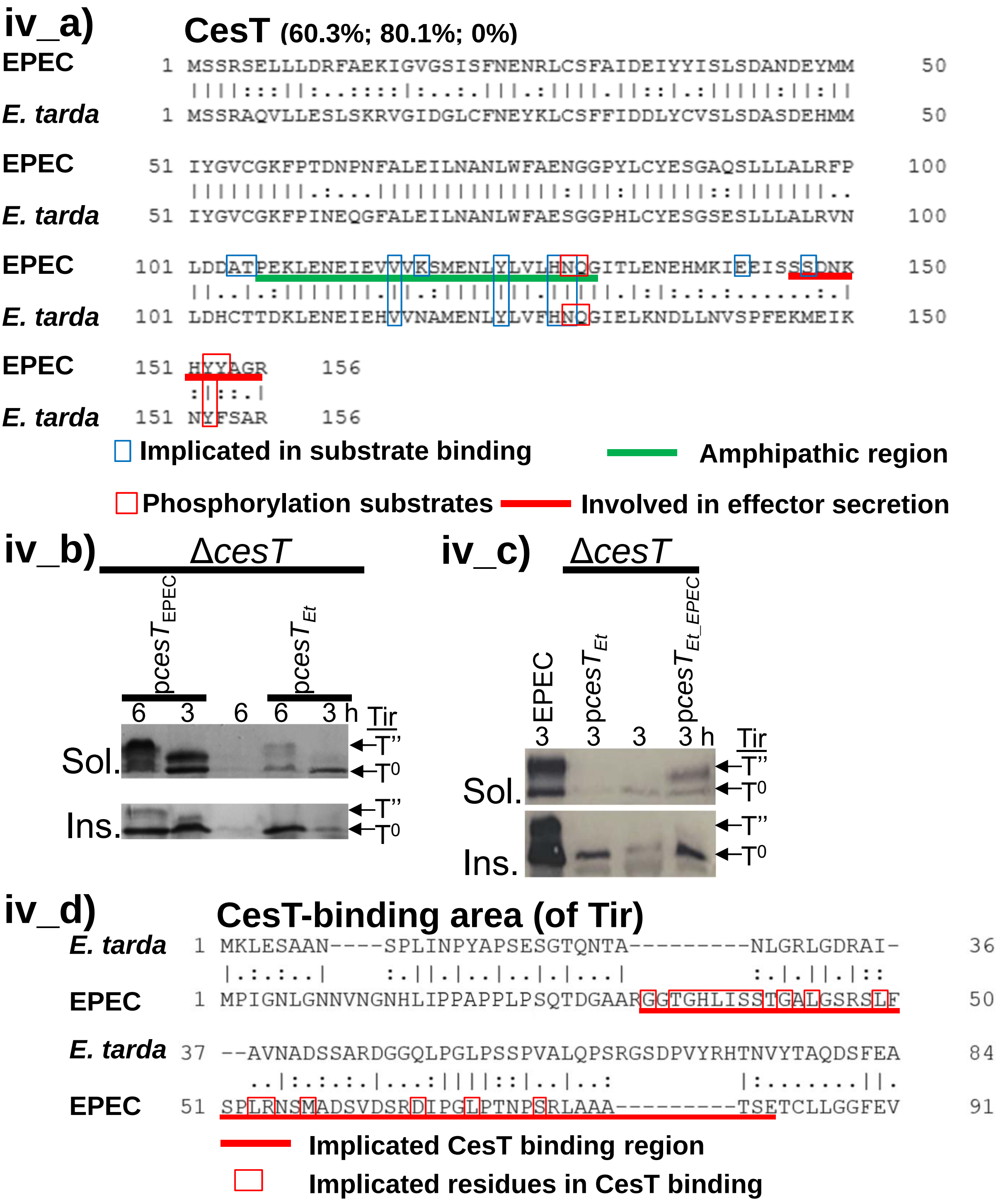

iv_a)
CesT (60.3%; 80.1%; 0%)
EPEC 1 MSSRSELLLDRFAEKIGVGSISFNENRLCSFAIDEIYYISLSDANDEYMM 50
E. tarda 1 MSSRAQVLLESLSKRVGIDGLCFNEYKLCSFFIDDLYCVSLSDASDEHMM 50
EPEC 51 IYGVCGKFPTDNPNFALEILNANLWFAENGGPYLCYESGAQSLLLALRFP 100
E. tarda 51 IYGVCGKFPINEQGFALEILNANLWFAESGGPHLCYESGSESLLLALRVN 100
EPEC 101 LDDATPEKLENEIEVVVKSMENLYLVLHNQGITLENEHMKIEEISSSDNK 150
E. tarda 101 LDHCTTDKLENEIEHVVNAMENLYLVFHNQGIELKNDLLNVSPFEKMEIK 150
EPEC 151 HYYAGR 156
E. tarda 151 NYFSAR 156
Implicated in substrate binding
Amphipathic region
Phosphorylation substrates
Involved in effector secretion
iv_b)
ΔcesT
pcesT EPEC
pcesT Et
6 3 6 6 3 h
Tir
Sol.
Ins.
T''
T0
iv_c)
ΔcesT
EPEC
pcesT Et
pcesT Et_EPEC
3 3 3 3 h
Tir
Sol.
Ins.
T''
T0
iv_d)
CesT-binding area (of Tir)
E. tarda 1 MKLESAAN----SPLINPYAPSESGTQNTA---------NLGRLGDRAI- 36
EPEC 1 MPIGNLGNNVNGNHLIPPAPPLPSQTDGAARGGTGHLISSTGALGSRSLF 50
E. tarda 37 --AVNADSSARDGGQLPGLPSSPVALQPSRGSDPVYRHTNVYTAQDSFEA 84
EPEC 51 SPLRNSMADSVDSRDIPGLPTNPSRLAAA---------TSETCLLGGFEV 91
Implicated CesT binding region
Implicated residues in CesT binding


### Figure S2

**v_a)** **EscP (21%; 39%; 25.4%)**

```
EPEC       1 MTRVSLKRNLIEPVSFRPSDDWPQGEESFNDIFYHIKKRSPDTLFIRMAS     50
                                                :||||.|...||.|.
E. tarda   1 -----------------------------------MKKRSADRWVIRGAE     15

EPEC      51 LWKRLFYSSGRRRRYFEEGEHSFSILCGRLRGVVLTIKCSNGIIYLSIKI    100
             ||..:|:|......:...|..:|.:....|:.:::.|:.:|...::::|:
E. tarda  16 LWFSMFHSGDNLFAHIPYGIRTFRLYNRVLKDIIIAIERNNQGGFITLKV     65

EPEC     101 SQNNRNHVFLYHKKDYVFDKLKEIFPDEAIEFTIEYEN    138
             |:|...|..:......:.:|:|.||...:|:|.|.|.:
E. tarda  66 SRNRIEHKVIIDNATIIIEKIKNIFDSCSIQFGITYYD    103
```

**v_b)**

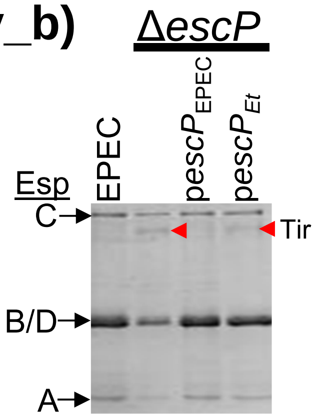


**v_c)**

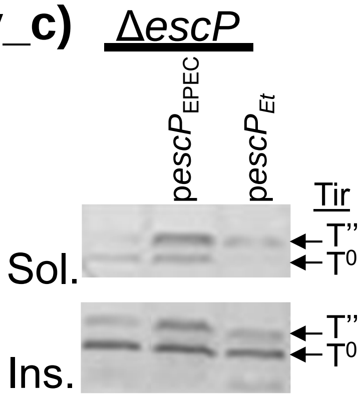


### Figure S2

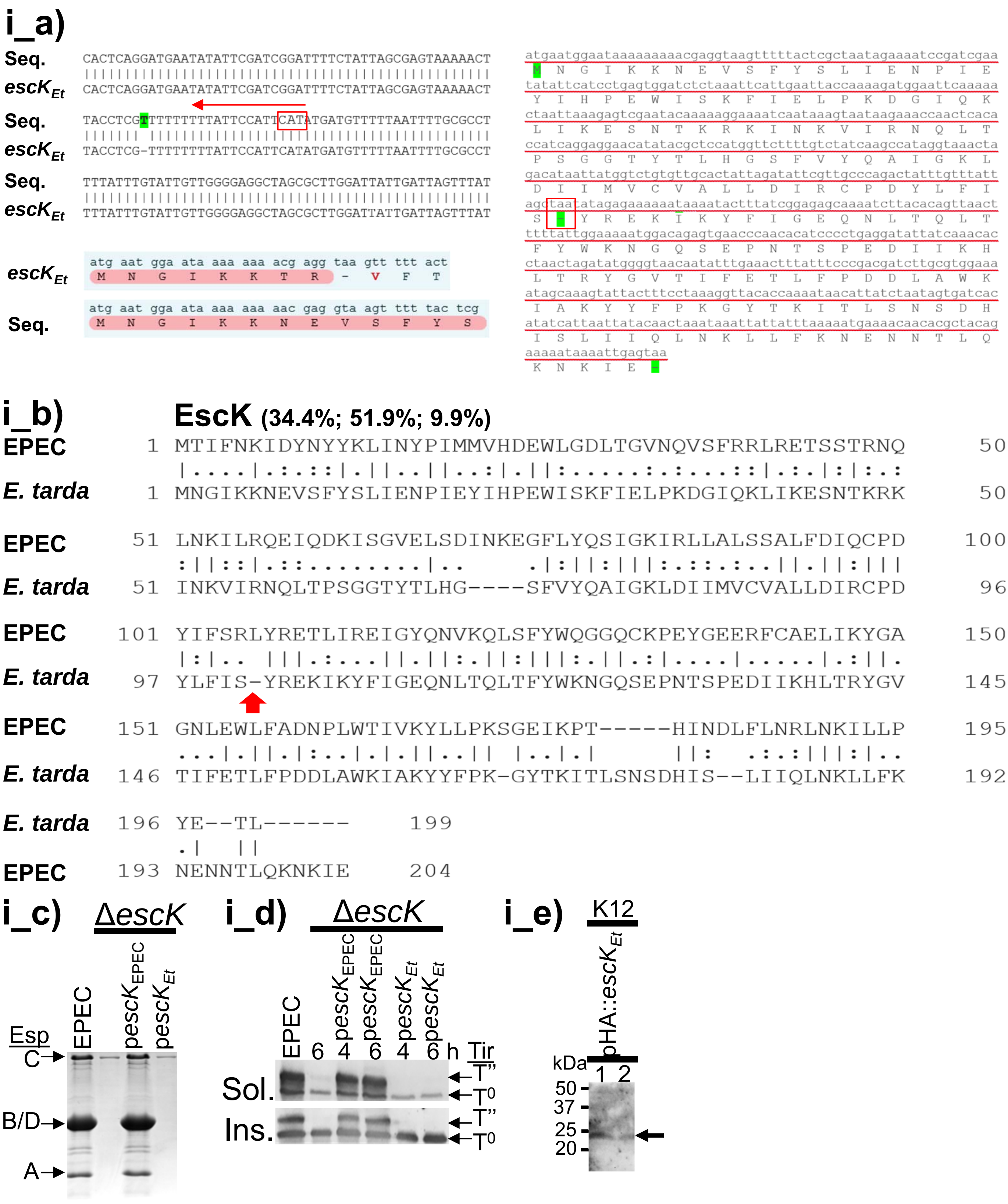
i_a)
Seq.
escK$_{Et}$
CACTCAGGATGAATATATTCGATCGGATTTTCTATTAGCGAGTAAAAACT
CACTCAGGATGAATATATTCGATCGGATTTTCTATTAGCGAGTAAAAACT
TACCTCGTTTTTTTTTATTCCATTCATATGATGTTTTTAATTTTGCGCCT
TACCTCG-TTTTTTTTTATTCCATTCATATGATGTTTTTAATTTTGCGCCT
TTTATTTGTATTGTTGGGGAGGCTAGCGCTTGGATTATTGATTAGTTTAT
TTTATTTGTATTGTTGGGGAGGCTAGCGCTTGGATTATTGATTAGTTTAT
atg aat gga ata aaa aaa acg agg taa gtt ttt act
M N G I K K T R - V F T
atg aat gga ata aaa aaa aac gag gta agt ttt tac tcg
M N G I K K N E V S F Y S
i_b)
EscK (34.4%; 51.9%; 9.9%)
EPEC 1 MTIFNKIDYNYYKLINYPIMMVHDEWLGDLTGVNQVSFRRLRETSSTRNQ 50
E. tarda 1 MNGIKKNEVSFYSLIENPIEYIHPEWISKFIELPKDGIQKLIKESNTKRK 50
EPEC 51 LNKILRQEIQDKISGVELSDINKEGFLYQSIGKIRLLALSSALFDIQCPD 100
E. tarda 51 INKVIRNQLTPSGGTYTLHG----SFVYQAIGKLDIIMVCVALLDIRCPD 96
EPEC 101 YIFSRLYRETLIREIGYQNVKQLSFYWQGGQCKPEYGEERFCAELIKYGA 150
E. tarda 97 YLFIS-YREKIKYFIGEQNLTQLTFYWKNGQSEPNTSPEDIIKHLTRYGV 145
EPEC 151 GNLEWLFADNPLWTIVKYLLPKSGEIKPT-----HINDLFLNRLNKILLP 195
E. tarda 146 TIFETLFPDDLAWKIAKYYFPK-GYTKITLSNSDHIS--LIIQLNKLLFK 192
E. tarda 196 YE--TL------ 199
EPEC 193 NENNTLQKNKIE 204
i_c)
ΔescK
EPEC
pescK$_{EPEC}$
pescK$_{Et}$
Esp
C
B/D
A
i_d)
ΔescK
EPEC
6
pescK$_{EPEC}$ 4
pescK$_{EPEC}$ 6
pescK$_{Et}$ 4
pescK$_{Et}$ 6 h
Tir
Sol.
Ins.
T"
T0
i_e)
K12
pHA::escK$_{Et}$
kDa
1 2
50
37
25
20

### Figure S3

## ii_a)

**SepD (39.6%; 63.6%; 4.1%)**

```
E. tarda    1 MSNSNDFS---IEWLSSLEFINNVSGDSSQIFFTIYKKDAILRDFGHSWI    47
              |:|:|..:   .:||::|:|:.:|:|..:.:.|.||:|:|.|.|||:.|:
EPEC        1 MNNNNGIAKNDCDWLTALDFVKDVNGSPTHLTFYIYQKNAFLHDFGNYWV    50

E. tarda   48 LYMKLNVDYKQMDTARLNKLCSILTAGKEYRHMGVYLSNNNWFLCRSYLK    97
              ||::|:.|::|:.|....:||:||...|||:.||::|||..|:||:.:.|
EPEC       51 LYIELSGDFRQIPTDTFIRLCNILAVSKEYKQMGIFLSNKKWYLCQIFHK   100

E. tarda   98 NGNHRSHISLIIMQHTFAVLLDKNFAKIDDEES--------HSHSSYSNI   139
              :.|||:::|..|||||.|.||||.|.|::...|        |..|....|
EPEC      101 DNNHRANMSKAIMQHTLASLLDKQFDKLEQSSSSDTMMPPTHLFSDIGRI   150

E. tarda  140 INLA      143
              :
EPEC      151 V---      151
```

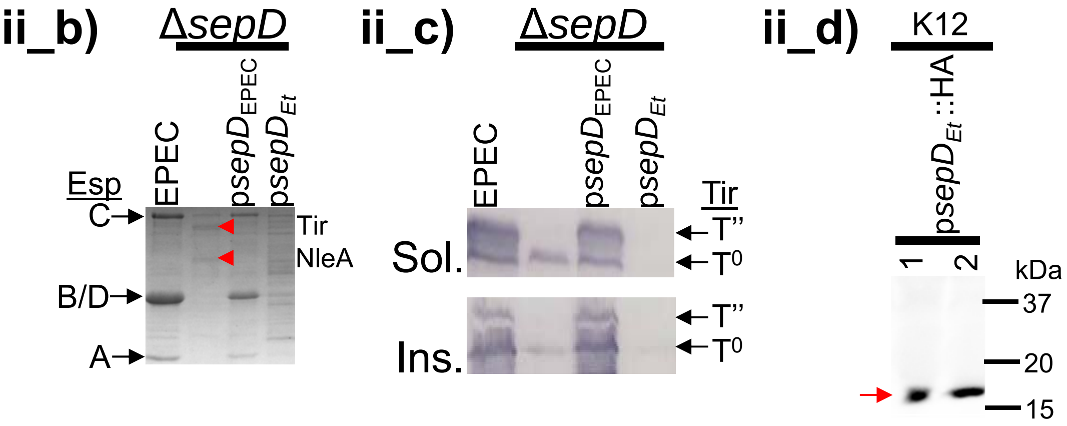


### Figure S3

#### iii_a) EscQ (21%; 44.1%; 8.9%)

```
E. tarda    1 ----------MIKLEYLSLFSKIGSGRVYHRDESGSIYLHFSHGAGRGVV    40
                        .|...||.|...||:||:|...|..:.:.:|...:|.|:.
EPEC        1 MKPLSSQLNMKINDFYLPLLPVIGTGRLYITSEGHACHAYFREVSGHGIR    50

E. tarda   41 IDLVVDSVPGQIWFSESEWLEWCNDIVHFSDIEAIPISLKDILATWVL--    88
              ..|......|:.|.||.::::||.::..:|:...||..:..::..||:
EPEC       51 FTLTYSGYEGRFWISEEQFIQWCQELFPYSESRLIPEDMIKLMILWVMQT   100

E. tarda   89 ----GDFFFGHKIIIKQITTLVLLDGFYPQATINSEDKNIAMVMLSTPLS   134
                  ||      :.:..:...:|....||....|:.:..:.:::|.|.:.
EPEC      101 ALPEGD------VSVDDVQFTMLNKDVYPVIENNNGENRLNVIILETTVQ   144

E. tarda  135 FLSFFGNGDSEIYPYGQFKLFDFEFYLAIGYDEIVISQLELGGAIRVNNI   184
              .|....|.:.:..|:.....||  .|:..|:.:..::|.:|.::|:.::
EPEC      145 SLQHLINDNWQFVPHSNALFFD--GYIVPGWTDYPVTELRVGDSLRLYHV   192

E. tarda  185 SNIDDDQVWISYGSPLGIVRVENSKLNIIETLYSPAAIVLDDAHDYEHMY   234
              .:..:.:.||...:||..|::.::.|::.:.|.:.....:........:
EPEC      193 DDSQERRCWIVINNPLATVKLSDNNLSVTDVLAADPLCTISSETVMNRI-   241

E. tarda  235 YYCV-GKVTLNISELDRFLVSGAVNYNKIMYFPNVKIINGKSCIAEGELV   283
               ||| |.|.::|..|.........:......|...::...:.||.|.::
EPEC      242 -YCVIGTVHIDIHMLRNVQKDDIIDSTGYHLFGGCQLVRNNTTIAYGSII   290

E. tarda  284 KINDAFYVELLNLE-    297
              |||:.||..:..:.
EPEC      291 KINEDFYFTISVVCD    305
```

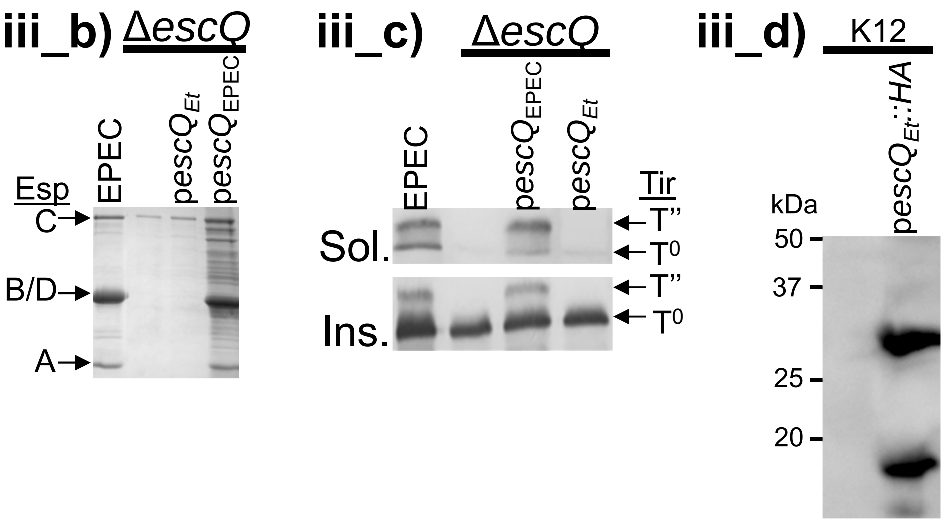


### Figure S3

**iv_a)** **EscN** **(62.1%; 76.4%; 4.3%)**

```
E. tarda    1 MI----SVLKFWGQLMDMNNEYVPKLCCENIKKLLRHDDVVISIENTATY    46
              ||    |||:.:.::..:.|..||.|                |:.::..|
EPEC        1 MISEHDSVLEKYPRIQKVLNSTVPAL----------------SLNSSTRY    34

E. tarda   47 EGKIIEIGAIFIKAYLPLAKIGSIYQIKPSDEFAEVISINEMSVTLLPFC    96
              |||||.||...|||.||.|:||:.|:|:||...||||:|:|..|.||||.
EPEC       35 EGKIINIGGTIIKARLPKARIGAFYKIEPSQRLAEVIAIDEDEVFLLPFE    84

E. tarda   97 DMSGMFLGQWLSFYQSEFFISVGYGLLGRVVNGLGHPLAAGDKLTNIPYQ   146
              .:|||:.|||||:...||.|.||..|||||:::|:|.|:.:......:|::
EPEC       85 HVSGMYCGQWLSYQGDEFKIRVGDALLGRLIDGIGRPMESNIVAPYLPFE   134

E. tarda  147 RSLFIEPPDPLSRAIIDKPLSVGVKAIDGLLTCGVGQRIGIFAGSGVGKS   196
              |||:.||||||.|.:||:|..:||:||||||||||:||||||||||||||
EPEC      135 RSLYAEPPDPLLRQVIDQPFILGVRAIDGLLTCGIGQRIGIFAGSGVGKS   184

E. tarda  197 TLLGMICNGAQADIIVMALIGERGREVNEFLALLPEETRLRSVFVVTTSD   246
              ||||||||||.|||||:|||||||||||||||||||:.|..:.|.||||||
EPEC      185 TLLGMICNGASADIIVLALIGERGREVNEFLALLPQSTLSKCVLVVTTSD   234

E. tarda  247 RPPLERMKAAFTATTIAEFFRDQGKHVLLIMDSVTRYARAARDVGLSIGE   296
              ||.||||||||||||||||:||||||:|||:||||||||||||||||:.||
EPEC      235 RPALERMKAAFTATTIAEYFRDQGKNVLLMMDSVTRYARAARDVGLASGE   284

E. tarda  297 PDIRGGFPPSVFATLPRLLERAGPAAVGAITAIYSVLIENDDANDPIADE   346
              ||:||||||||||::||:||||||||..|:||||||:||:|:|:.||||.||
EPEC      285 PDVRGGFPPSVFSSLPKLLERAGPAPKGSITAIYTVLLESDNVNDPIGDE   334

E. tarda  347 VRSILDGHIILSRKLAEENHYPAIDVGLSASRIMINVTSREQSNAAATLK   396
              |||||||||:|:|:|||||||:||||:||||||:|||||:|.||.:.|...|||..|
EPEC      335 VRSILDGHIVLTRELAEENHFPAIDIGLSASRVMHNVVTSEHLRAAAECK   384

E. tarda  397 SMIATYKDVELLLRIGEYKHGEDPQVDRAIHLWPQIQSFCRQPFDSVMEF   446
              .:|||||:||||:|||||..|:||:.|:||.....||:|.:|....:..:
EPEC      385 KLIATYKNVELLIRIGEYTMGQDPEADKAIKNRKLIQNFIQQSTKDISSY   434

E. tarda  447 NTTINELFRTVG     458
              ..||..||:.|.
EPEC      435 EKTIESLFKVVA     446
```

**iv_b)**

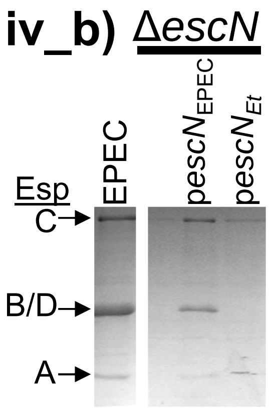


**iv_c)**

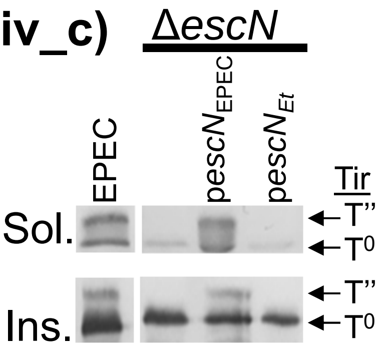


**iv_d)**

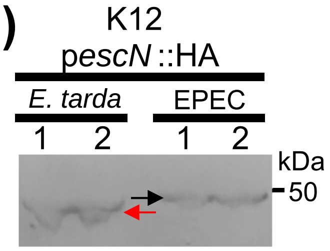


### Figure S3

#### v_a) **EscV** (66.3%; 79.9%; 1.6%)

```
E. tarda    1 MPLKMKSYISFLNSFSKYQDLVLAIFFFMAVMMMIIPLPTIVVDIIIAIN    50
                  |...::.......|.||:||:||||||||||||||||:|||||||||
EPEC        1 ----MNKLLNIFKKAESYHDLILALFFFMAVMMMIIPLPTVVVDIIIAIN    46

E. tarda   51 ISTALLLLMLSIYIKRPLDLSVFPTILLITTLMRLSLSVSTTRLILLQHD   100
              ||||||||||||||.||:|:.|||||||||||||||||||||||||.||
EPEC       47 ISTALLLLMLSIYIKNPLELTSFPTILLITTLMRLSLSVSTTRLILLHHD    96

E. tarda  101 AGEIIYSFGNFVVGGNIVVGLIIFVIITIVQFLVITKGAERVAEVGARFS   150
              ||:||||||||||||||||||:||.|||||||:|||||||||||||.||||
EPEC       97 AGDIIYSFGNFVVGGNIVVGLVIFTIITIVQFMVITKGAERVAEVSARFS   146

E. tarda  151 LDGMPGKQMSIDGDMRAGIIDAQEAKKQRAEVQKESQLYGAMDGAMKFVK   200
              ||||||||||||||||||:||..|||..|:.||||||.||:|||||||||
EPEC      147 LDGMPGKQMSIDGDMRAGVIDPLEAKVLRSRVQKESQFYGSMDGAMKFVK   196

E. tarda  201 GDAIASIIIIVVNLLGGVLIGMWQFNMSFSDALALFSVLSVGDALVAQIP   250
              |||||.|||::|||.||||||||||||:|.||:||:|||||||||||||||||
EPEC      197 GDAIAGIIIVLVNLFGGVLIGMWQFDMPFSEALSLFSVLSVGDALVAQIP   246

E. tarda  251 ALLISVTAGIVVTRVPGEI--NNNLAGDIIGQVSGNSRPFFISAGLMLVM   298
              ||:||||||:||||||||.  ..||||||:.|||.|||||.|||.|||||
EPEC      247 ALIISVTAGVVVTRVPGESEKEENLAGDIVQQVSVNSRPFLISAALMLVM   296
```

**C-terminal domain (58.3%; 75.8%; 0.3%)**

```
E. tarda  299 ASIPGFPVLVFILLSLAMGVIGWRVNKKQVF---DSHSSLGPEERIGSEN   345
              |.|||||.|||:.|::.:..|.|::.||:.|   ::..::| .:...|:|
EPEC      297 AIIPGFPTLVFLFLAVCLLGIAWKLQKKRTFGAGNNKDAMG-ADLSNSQN   345

E. tarda  346 VTSGAVPLRLKIGARIFNAELKQFIDMMRWRFFDERGIPLPEIVLECSSI   395
              ::.||.||.|.:.:.|::::.|.|::|||.||:|.|||||:|::.....
EPEC      346 ISPGAEPLILNLSSNIYSSDITQQIEVMRWNFFEESGIPLPKIIVNPVKN   395
```

α

```
E. tarda  396 KDDAIEVYLYQELVYKTFWENEKIIFFPNSNDFSFEYQKELLPYGRKVYK   445
              .|.|||..||||.:||....::.:.|.....:.|||:.:|.|.....|||
EPEC      396 NDSAIEFLLYQESIYKDTLIDDTVYFEAGHAEISFEFVQEKLSTNSIVYK   445
```

β

```
E. tarda  446 TPLANEEIAISLGIKVVNNVNDAVVFYSKLIVSCCAREFIGVQETRYLMD   495
              |...|:::|...|:.|....||.:.|..|.:|...|:|||||||||||||
EPEC      446 TNKTNQQLAHLTGMDVYATTNDKITFLLKKLVLSNAKEFIGVQETRYLMD   495
```

Y

```
E. tarda  496 IMERQYSELVKELQRQLGLSKIVDILQRLVDEEISIRDLRTIFETLILWS   545
              ||||:|:||||||||||||||||||||||||:|.:||||||||||||||.||
EPEC      496 IMERKYNELVKELQRQLGLSKIVDILQRLVEENVSIRDLRTIFETLIFWS   545

E. tarda  546 VKEKDVVILCEYVRVALRRHIISKHSNHGEMLSVWLIGSELENEMREAIR   595
              .|||||||||||||:||||||:.::|..|.:|:||||||::||||:|||:||
EPEC      546 TKEKDVVILCEYVRIALRRHILGRYSVSGTLLNVWLIGSDIENELRESIR   595

E. tarda  596 QTSSGSYLNLSAERSSQLIASMNTIIDSNINGVILTSLDIRRYMKKLIEG   645
              |||||||||:|.||:.|:|..:..|::...||||||:|||||||:||:|||
EPEC      596 QTSSGSYLNISPERTEQIIGFLKNIMNPTGNGVILTALDIRRYVKKMIEG   645

E. tarda  646 TIPTISVLSFQEVGNDIGLKVLGTINGYE-    674
              :.|::.|||||||||:|.||||||:|.:.
EPEC      646 SFPSVPVLSFQEVGNNIELKVLGTVNDFRA    675
```

αβY= SepL binding sites

### Figure S3

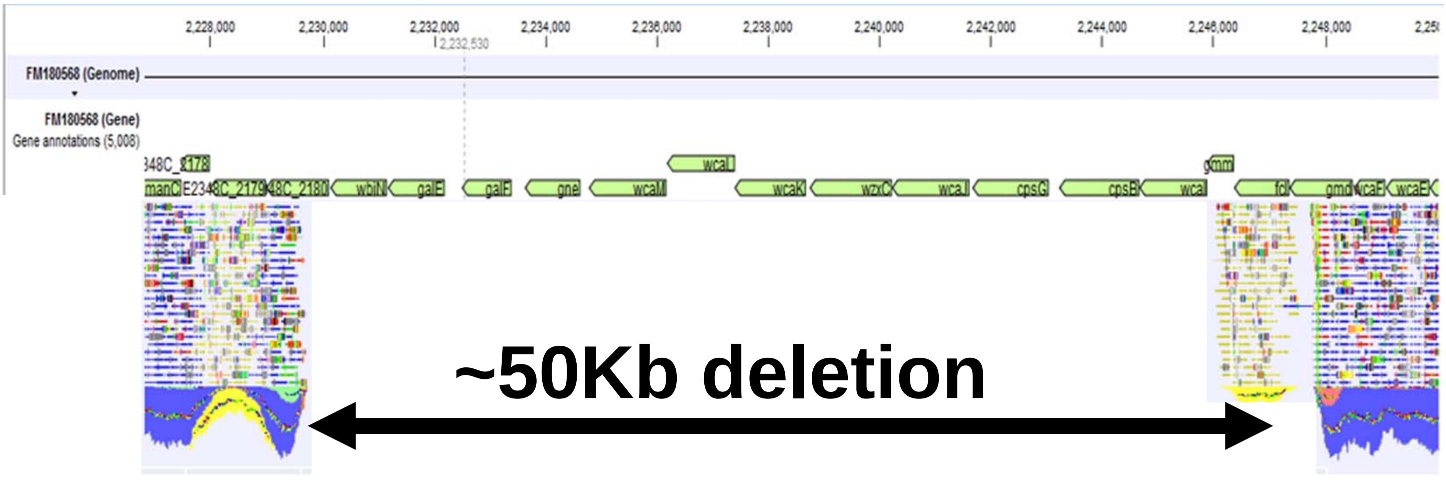


| | | | | | |
|---|---|---|---|---|---|
| CDS | 2227497 | 2227997 - | E2348_C_2184 | gmm | GDP-mannose mannosyl hydrolase |
| CDS | 2228000 | 2228965 - | E2348_C_2185 | fcl | bifunctional GDP-fucose synthetase:GDP-4-dehydro-6-deoxy-D-mannose epimerase/GDP-4-dehydro-6-L-deoxygalactose reductase |
| CDS | 2228968 | 2230089 - | E2348_C_2186 | gmd | GDP-D-mannose dehydratase, NAD(P)-binding |
| CDS | 2230116 | 2231132 - | E2348_C_2187 | wbiN | predicted glycosyltransferase |
| CDS | 2231151 | 2232170 - | E2348_C_2188 | galE | UDP-galactose-4-epimerase |
| CDS | 2232480 | 2233373 - | E2348_C_2189 | galF | predicted subunit with GalU |
| CDS | 2233616 | 2234611 - | E2348_C_2190 | gne | putative UDP-galactose 4-epimerase |
| CDS | 2234769 | 2236163 - | E2348_C_2191 | wcaM | predicted colanic acid biosynthesis protein |
| CDS | 2236174 | 2237394 - | E2348_C_2192 | wcaL | predicted glycosyl transferase |
| CDS | 2237391 | 2238671 - | E2348_C_2193 | wcaK | predicted pyruvyl transferase WcaK |
| CDS | 2238743 | 2240221 - | E2348_C_2194 | wzxC | colanic acid exporter |
| CDS | 2240223 | 2241617 - | E2348_C_2195 | wcaJ | predicted UDP-glucose lipid carrier transferase |
| CDS | 2241672 | 2243042 - | E2348_C_2196 | cpsG | phosphomannomutase |
| CDS | 2243235 | 2244671 - | E2348_C_2197 | cpsB | mannose-1-phosphate guanyltransferase |
| CDS | 2244674 | 2245897 - | E2348_C_2198 | wcaI | predicted glycosyl transferase |
| CDS | 2245894 | 2246373 - | E2348_C_2199 | wcaH | GDP-mannose mannosyl hydrolase |
| CDS | 2246376 | 2247341 - | E2348_C_2200 | wcaG | bifunctional GDP-fucose synthetase:GDP-4-dehydro-6-deoxy-D-mannose epimerase/GDP-4-dehydro-6-L-deoxygalactose reductase |
| CDS | 2247344 | 2248465 - | E2348_C_2201 | gmd | GDP-D-mannose dehydratase, NAD(P)-binding |
| CDS | 2248492 | 2249040 - | E2348_C_2202 | wcaF | predicted acyl transferase |
| CDS | 2249056 | 2249802 - | E2348_C_2203 | wcaE | predicted glycosyl transferase |

O127-antigen Biosynthesis Locus

Colanic Acid Production

### Figure S4

```
EscP_Salm      MRILNQDDFNYYKLINHPLT-VIHSDWITELTGVSRLCYRNLIENNATRSQLNNVLKMKF    59
EscP_EPEC      MT-----RVSLKRNLIEPVSFRPSDDWPQGEESFNDIF---------------------Y    34
EscP_tarda     ------------------------------------------------------------    0


EscP_Salm      QLITESMEDFFVDKNKLVYQIIGKAKIMAISGALFEMKCPDYLFSGHYRDHLINELGYEN    119
EscP_EPEC      HIKKRSPDTLFIRMASLW------------------------------------------    52
EscP_tarda     -MKKRSADRWVIRGAELW------------------------------------------    17
                : ..* :  .:   .*

EscP_Salm      VKQLSFFWKGGDGRAEYTNTNFCDKLLAYGSGNLEYIFRNEPLWEIVKYLL-PKGG----    174
EscP_EPEC      ---KRLFYSSGRRRRYFE----------EGEHS--FSILCGRLRGVVLTIKCSNGIIYLS    97
EscP_tarda     ---FSMFHSGDNLFAHIP----------YGIRT--FRLYNRVLKDIIIAIERNNQGGFIT    62
                    :* ...                   *  .  : :    *  ::  :   :

EscP_Salm      -EIKANNIDENFLNR--------LNRILSPYEAL--------      199
EscP_EPEC      IKISQNNRNHVFLYHKKDYVFDKLKEIF-PDEAIEFTIEYEN      138
EscP_tarda     LKVSRNRIEHKVIIDNATIIIEKIKNIF-DSCSIQFGITYYD      103
                ::. *. :. .:          ::.*:    ::
---------------------------------------------------------------------------
EscI_tarda     -----------------MDPILQTNALSSILPHSAEASTIPSSVQLTNMPA-QPVEMTAA    42
EscI_EPEC      MDALCYCLSHEKRLTVNMNNINQSENINIQLNKASQTNVV-----DEHIPLASTPSAAGA    55
EscI_Salm      -----------------MSFIQ---NIN-----EMTTNVINK---EKGESLQPSPSAESA    32
                                *. *     :.        :..:               .  .*

EscI_tarda     SQFLSQLLPSTASVASPEQVLIEEIKRRHFNALNAPDLNFDDLAAGKLSPTDMLQLQRSV    102
EscI_EPEC      AQFLDQLLPKTAGVSSPEQVLIEEIKKRHLATMN-TDLSFDALSAGGLSPEDVLTLQKNV    114
EscI_Salm      AQFLDQLLPKTAAIASPEQVLIEEIKKRHLTTMN-TNLSLDTLSAGGLSPENLIDLQKTV    91
               :***.****.**.::***********:**: ::*  :*.:* *:** *** ::: **:.*

EscI_tarda     LNANVNIDVISKLASLFSTSITKLTSMQ    130
EscI_EPEC      LNANVNVDVVSKLASLLSTSVTKLVSMQ    142
EscI_Salm      LNANINIDVVSKFASLLSTSITKLVSVQ    119
               ****:*:**:**:***:***:***.*:*
---------------------------------------------------------------------------
EscE_Salm      MMSLTELEDMLIEDTENNSEKLLSILKERRKQLNKSLENPLDKLAYKKTEQMLVACEAAQ    60
EscE_EPEC      MITITELEDEIIKNKEAANVFI-EKINDKKNEIHEKMKHPLDKVTYNEAKELLIACDAAI    59
EscE_tarda     MISITLLEDIIIEDPEQSTSELIKTLEYEIEKNNHEMKMIKCNAEYKKRDKLNIAYLSAI    60
               *:::* *** :*:: *  .  : . :: . :: :..::    :  *:: .:: :*  :*

EscE_Salm      QVIELIAFRFSNNKKKGDIHEKKN        84
EscE_EPEC      RTIEIMRIRINNK-----------        72
EscE_tarda     STITTIKSRIKNEKNR--------        76
                .*  :  *:.*:
```

### Figure S5
